# A mitochondrial orphan protein contributes to sperm development in bivalves

**DOI:** 10.64898/2026.11.03.763971

**Authors:** Julie Brémaud, Laurie Touchette, Marianne Hubert, Annie Angers, Sophie Breton

## Abstract

Lineage-specific genes, or orphan genes, are increasingly recognized as important drivers of evolutionary innovation, yet their functional characterization remains largely restricted to nuclear genomes of model organisms. Whether lineage-specific genes encoded by mitochondrial DNA contribute to phenotypic diversification remains virtually unexplored. Here we investigate MORF, a rapidly evolving ORFan protein encoded by the paternally transmitted mitochondrial genome of bivalves with doubly uniparental inheritance (DUI). Combining immunolocalization, functional neutralization, protein interaction assays and proteomics in the distantly related bivalves *Mytilus edulis* and *Venustaconcha ellipsiformis*, we characterize MORF localization, function and molecular associations during male germ cell development. MORF localizes to developing sperm cells and mature sperm mitochondria in both species and is additionally detected in the acrosome of *M. edulis*. In cultured *M. edulis* germ cells, antibody-mediated neutralization of MORF increases the frequency of spermatids displaying four rather than the canonical five mitochondria, providing functional evidence that MORF contributes to mitochondrial organization during spermatogenesis. Comparative proteomic analyses identify MORF-associated proteins involved in cytoskeletal organization, protein homeostasis and extracellular matrix organization. The association between MORF and laminin is independently supported by western blotting in both species. Despite extensive sequence divergence, the two MORF proteins display overlapping functional and molecular features. Together, these findings uncover a functional role for a rapidly evolving mitochondrial orphan protein and point to mitochondrial orphan genes as a potentially overlooked source of evolutionary novelty.

## INTRODUCTION

How newly emerged genes acquire biological functions and become integrated into pre-existing cellular networks remains one of the central questions in evolutionary biology. Lineage-specific genes, also known as taxonomically restricted or orphan genes, which lack detectable homologs outside a species or evolutionary lineage, provide a unique opportunity to address this question. Once considered, and still sometimes dismissed, as rare evolutionary curiosities or annotation artifacts, these genes are now increasingly recognized as important drivers of lineage-specific adaptations and phenotypic innovation (1–3).

Over recent years, functional studies have demonstrated that lineage-specific genes participate in diverse biological processes, including development, morphogenesis, neuronal maturation, stress tolerance, reproduction and sexuality (2–4). However, the mechanisms through which they acquire and maintain biological functions remain poorly understood, as functional evidence is largely restricted to lineage-specific genes from nuclear genomes in a limited number of model organisms (2–4). Comparable questions have received far less attention in mitochondrial genomes. Although animal mitogenomes are often portrayed as compact and evolutionarily constrained, encoding only thirteen oxidative phosphorylation proteins together with two ribosomal and 22 transfer RNAs (5), growing evidence reveals unexpected diversity in their organization and gene content, including lineage-specific open reading frames identified in several animal lineages (5–8). Yet, the biological roles of these mitochondrial lineage-specific genes remain virtually unexplored, and their potential contribution to phenotypic evolution is unknown. Establishing whether mitochondrial lineage-specific genes can acquire adaptive functions therefore represents an important challenge for understanding the evolutionary potential of mitochondrial genomes beyond bioenergetics.

Bivalve mollusks exhibiting doubly uniparental inheritance (DUI) of mitochondria provide a unique natural system to address this question. Unlike the strict maternal inheritance of mitochondrial DNA (mtDNA) found in nearly all animals, DUI maintains distinct maternal (F-type) and paternal (M-type) mitogenomes that are transmitted according to sex (9–11). This remarkable system has persisted independently across deeply divergent bivalve clades for more than 200 million years, yet its molecular basis remains poorly understood (12, 13). One distinctive feature of DUI mitogenomes is the presence of lineage-specific mitochondrial genes. In freshwater mussels (order Unionida), a female-specific *forf* and male-specific *morf* genes are tightly associated with DUI and gonochorism, whereas hermaphroditic species exhibiting strict maternal inheritance instead possess a highly modified *horf* gene replacing *forf* (14). F- and M-specific mitochondrial orphan genes have also been identified in marine mussels (Mytilida) and the marine clam *Ruditapes philippinarum* (Venerida), despite extreme sequence divergence that obscures their evolutionary relationships (15). Regardless of their evolutionary origin, their strict association with sex across independently evolved DUI lineages strongly suggests that they perform specialized functions related to reproductive biology, sex determination or the transmission of paternal mitochondria (12–15).

The ORFan protein MORF, encoded by the *morf* gene, is a strong candidate for mediating male-specific functions. A recent study has shown that MORF expression is restricted to male gonads during the reproductive season in both marine and freshwater mussels, whereas FORF exhibits a broader expression profile (16). Immunolocalization in the marine mussel *Mytilus edulis* further revealed MORF within sperm mitochondria as well as in the acrosomes of late spermatids and mature spermatozoa, suggesting male-specific functions extending beyond mitochondrial metabolism (16, 17). Structural predictions further suggest interactions with cytoskeletal components, ubiquitination pathways, and apoptosis-related processes, leading to the hypothesis that MORF could participate in the selective retention of paternal mitochondria during embryogenesis (15). However, these hypotheses remain largely speculative, as no comprehensive characterization of MORF molecular interactions has been performed. Moreover, MORF proteins exhibit extraordinary sequence divergence across bivalve lineages, raising a fundamental evolutionary question: do these rapidly evolving lineage-specific proteins retain conserved biological functions despite extensive primary sequence divergence, or have they evolved species-specific molecular mechanisms?

Here, we investigate the molecular function of MORF in two distantly related bivalves representing independent DUI lineages, the marine mussel *Mytilus edulis* (Mytilida) and the freshwater mussel *Venustaconcha ellipsiformis* (Unionida). By integrating AlphaFold structural predictions, immunohistochemical localization, co-immunoprecipitation, GST pull-down proteomics and biochemical validation, we characterize MORF from protein structure to interaction networks and cellular distribution. Despite extensive sequence divergence between species, we identify conserved interaction signatures centered on extracellular matrix components, particularly laminins, supporting an evolutionarily conserved role in spermatogenesis and sperm differentiation. Consistent with this hypothesis, antibody-mediated neutralization of MORF disrupts the canonical mitochondrial organization of developing spermatids, providing functional evidence for its role during sperm development. Together, these findings demonstrate that lineage-specific genes encoded by mitochondrial genomes can evolve conserved biological functions and contribute to the emergence of lineage-specific reproductive innovations, thereby expanding the evolutionary significance of mitochondrial genomes far beyond their canonical role in cellular energy metabolism.

## MATERIAL AND METHODS

### Sample collection and preparation

Adult *Mytilus edulis* were obtained from local seafood markets in Montreal, Canada, in June and July 2023 and in April 2024 (Prince Edward Island Aquafarm Inc., Canada). Adult *Venustaconcha ellipsiformis* were collected from the Meramec River (Missouri, USA) in October 2024 by the Missouri Department of Conservation and shipped alive on ice to the Université de Montréal. Sex was determined by microscopic examination of fresh gonadal smears, and only males exhibiting motile spermatozoa were retained for subsequent analyses.

For histological and immunohistological analyses, male gonads were dissected, embedded in OCT compound (Optimum Cutting Temperature; Sigma-Aldrich), snap-frozen in isopentane chilled with dry ice, and stored at −80 °C. Frozen samples were shipped on dry ice to the histology platform of the Institute for Research in Immunology and Cancer (IRIC, Université de Montréal), where they were cryosectioned (10 µm) using a Leica CM3050SA cryostat. Sections were either stained with hematoxylin and eosin (HE) for histological examination or stored at −20 °C until immunohistochemistry. For co-immunoprecipitations and pull-down assays, male gonads were dissected and stored at −80 °C until protein extraction.

### Immunohistochemistry

Cryosections were thawed at room temperature (RT), fixed in 4% paraformaldehyde (PFA) for 15 min, washed with distilled water, and rehydrated in phosphate-buffered saline (PBS). Sections were permeabilized with 1% Triton X-100 in PBS for 30 min at RT and washed in PBS. A hydrophobic barrier was drawn around each section, and non-specific binding was blocked with 4% bovine serum albumin (BSA) and 5% normal goat serum for 1 h at RT.

Sections were incubated overnight at 4 °C with rabbit anti-MORF antibodies (1:400 dilution for *M. edulis*; 1:800 dilution for *V. ellipsiformis*), mouse anti-ATP5A (Abcam, ab147448; 1:300), or rabbit anti-laminin (Invitrogen, PA1-16730; 1:500), all diluted in 1% BSA. The specificity of both anti-MORF antibodies has been previously validated (16, 17). Sections incubated with 1% BSA alone served as negative controls.

Following PBS washes, sections were incubated for 1 hour at RT with Alexa Fluor 488-goat anti-rabbit IgG (1:400; Thermo Fisher Scientific) or Alexa Fluor 594-goat anti-mouse IgG (4 µg/ml; Thermo Fisher Scientific). After additional washes, slides were mounted with DAPI-containing mounting medium (Abcam) and imaged using a Leica Mica fluorescence microscope equipped with Leica LAS X software.

### Culture of *Mytilus edulis* male germ cells

Primary cultures of *M. edulis* male germ cells were established using a protocol previously developed in our laboratory for mantle cell cultures that support male germ cell proliferation (58). Briefly, mature gonads were rinsed in seawater (Instant Ocean Sea Salt) and placed in artificial seawater (200 mM NaCl, 11 mM CaCl₂, 10 mM KCl, 27 mM MgCl₂, 27 mM MgSO₄, 2 mM NaHCO₃ and dextrose 6 g/l) supplemented with penicillin (100 U/ml) and streptomycin (100 µg/ml), minced, and enzymatically dissociated with 0.25% trypsin. Digestion was stopped by adding Leibovitz’s L-15 medium prepared in artificial seawater and supplemented with 20% newborn calf serum, penicillin (100 U/ml), and streptomycin (100 µg/ml), pH 7.4.

Cell suspensions were filtered through 100- and 40-µm cell strainers, washed by centrifugation, and resuspended in fresh medium. Cells were seeded onto poly-D-Lysine-coated six-well plates at a density of approximately 2 × 10⁷ cells per well, centrifuged briefly to promote attachment, and maintained at 25 °C. Cell adhesion and settling were assessed after 2 d of culture.

To determine whether MORF was released into the extracellular environment, conditioned culture media were collected after 2 d, cleared by centrifugation (14,000 rpm, 20 min), and the supernatants were retained for western blot analysis. Adhered cells were washed twice with ice-cold PBS and lysed in homogenization buffer (0.25 M sucrose, 1 mM EDTA, 10 mM HEPES, pH 7.4). Cells were collected by scraping, sonicated (3 × 3 s), and centrifuged (14,000 rpm, 15 min) to remove insoluble material. Supernatants were retained on ice. Protein concentrations in both conditioned media and cell lysates were determined by spectrophotometry before western blotting.

### MORF neutralization in cultured male germ cells

To investigate the putative role of MORF in male germ cell mitochondria organization, primary cultures of *M. edulis* male germ cells were established as described above. Germ cells isolated from three adult males were seeded onto poly-D-lysine-coated chamber slides in 24-well plates (1 ml culture medium per well). After 2 d at 25 °C, once cell adhesion had been confirmed, cultures were treated with one of three conditions: species-specific anti-MORF antibodies (20 μg/ml), rabbit pre-immune serum (20 μg/ml = control #1), or horseradish peroxidase (HRP)-conjugated goat anti-rabbit IgG (20 μg/ml = secondary antibody-only control #2 or Ab2). The latter two treatments served as negative controls. Each treatment was performed in triplicate, and cultures were maintained for an additional 5 d at 25 °C.

Following treatment, cells were washed with PBS and incubated with 500 nM MitoTracker Red CMXRos (Invitrogen) for 30 min at 25 °C to visualize mitochondria. Cells were then washed, fixed in 4% PFA, mounted in DAPI-containing mounting medium (Abcam), and imaged using a Leica Mica fluorescence microscope equipped with Leica LAS X software.

### Quantification of mitochondrial organization and statistical analyses

For each technical replicate, 20 microscopic fields were randomly acquired using identical imaging settings. The total number of spermatids was manually counted regardless of their orientation. Spermatids displaying four or five mitochondria were scored only when the mitochondrial rosette was entirely visible, corresponding to a transverse orientation of the cell (19). Cells in which the rosette could not be fully resolved (e.g., longitudinally oriented spermatids) were included in the total spermatid count but were excluded from the classification of four- or five-mitochondria rosettes. Free four- and five-mitochondria rosette structures were similarly counted only when all mitochondria composing the rosette were clearly visible.

Statistical analyses were performed in R (v4.5.2) using the lme4 and emmeans packages. The effect of MORF neutralization on mitochondrial organization was assessed using generalized linear mixed-effects models (GLMMs) with a binomial error distribution and logit link function. Treatment (pre-immune serum, secondary antibody, or anti-MORF antibody) was included as a fixed effect, whereas individual male identity was treated as a random effect. Pairwise comparisons were performed using estimated marginal means (EMMs) with Tukey’s adjustment for multiple testing. Because both control treatments produced comparable responses, they were pooled and compared with the anti-MORF treatment using Fisher’s exact test. Statistical significance was set at *P* < 0.05.

### Western blotting

Protein extracts were prepared in homogenization buffer (0.25 M sucrose, 1 mM EDTA, 10 mM HEPES, pH 7.4) supplemented with protease inhibitors (benzamidine, PMSF, leupeptin, and aprotinin), and sonicated. Total protein concentration was determined using the Bradford assay.

Protein samples were mixed with Laemmli buffer, heated at 95 °C for 5 min, and separated by tripartite tricine SDS-PAGE (59). Equal amounts of protein (100 μg per lane) were loaded for each sample. Proteins were transferred onto 0.45-μm nitrocellulose membranes, which were blocked in 5% non-fat dry milk before overnight incubation at 4 °C with species-specific anti-MORF antibodies (1:2,000) or rabbit anti-laminin (Invitrogen, PA1-16730; 1:5,000).

After washing in Tris-buffered saline containing 0.05% Tween-20 (TBS-T), membranes were incubated for 30 min at RT with HRP-conjugated goat anti-rabbit IgG (1:2,000). Immunoreactive proteins were detected using SuperSignal™ West Pico PLUS chemiluminescent substrate (Thermo Fisher Scientific) and imaged with a Fusion FX imaging system (Vilber Lourmat).

### Co-immunoprecipitation

Gonad protein extracts were prepared as described above. For each species, two independent biological replicates were analyzed. Protein A magnetic Sepharose beads (Cytiva) were washed and equilibrated in TBS (50 mM Tris-HCl, 150 mM NaCl, pH 7.5) supplemented with 2 M urea. To reduce non-specific binding, 3-4 mg of total protein were pre-cleared by incubation with washed beads for 2 h at RT.

Species-specific anti-MORF antibodies (25 μl, 0.5-1 mg/ml) or rabbit pre-immune serum (as negative control) were coupled to Protein A beads and covalently cross-linked with dimethyl pimelimidate in triethanolamine buffer. Residual reactive groups were quenched with ethanolamine before the beads were extensively washed.

Pre-cleared lysates were incubated with antibody-coupled beads for 1 h at RT. Following immunoprecipitation, the non-bound fraction was collected, and the beads were washed extensively with TBS containing 2 M urea. Immunoprecipitation efficiency was verified by western blotting using half of the recovered beads, whereas the remaining beads were submitted to the proteomics core facility of the Institute for Research in Immunology and Cancer (IRIC, Université de Montréal) for mass spectrometry analysis.

For western blot validation, bound proteins were eluted with 0.1 M glycine containing 2 M urea (pH 2.9). Eluted fractions, together with total homogenates and non-bound fractions, were analyzed by western blotting to confirm MORF immunoprecipitation.

### Pull-down assays

GST pull-down assays were performed using two independent biological replicates for each species. Competent *Escherichia coli* BL21(DE3) cells were transformed with expression plasmids encoding GST-tagged MORF from *M. edulis* or *V. ellipsiformis*, or with a GST-only expression vector. Recombinant proteins were expressed in LB medium supplemented with ampicillin and induced at mid-log phase (OD_600_ = 0.5-0.6) with isopropyl β-D-1-thiogalactopyranoside (IPTG) for 2.5 h at 37 °C.

Bacterial pellets were collected, lysed by sonication in PBS containing 8 M urea, supplemented with Triton X-100, and clarified by centrifugation. Bacterial lysates were pre-cleared with washed glutathione magnetic agarose beads (Pierce, Thermo Scientific) for 2 h at RT. Pre-cleared lysates were incubated with fresh glutathione beads for 3 h at RT under constant agitation. Following extensive washing, GST-MORF or GST-coupled beads were incubated with gonad protein extracts overnight at 4 °C. After washing, samples were resuspended in 100 mM Tris-HCl (pH 8). Bead-bound protein complexes were maintained on ice and submitted to the proteomics core facility at the Institute for Research in Immunology and Cancer (IRIC, Université de Montréal) for mass spectrometry analysis.

### Mass spectrometry

Mass spectrometric analysis of immunoprecipitated and pulled-down complexes were carried as follow. Protein samples were reconstituted in 50 mM ammonium bicarbonate containing 10 mM Tris(2-carboxyethyl)phosphine hydrochloride (TECP), alkylated with chloroacetamide, digested with trypsin, and resuspended in 5% ACN-4% formic acid (FA) prior to analysis. Peptides were separated by reverse-phase nano-liquid chromatography on an Easy nLC-1200 system (Thermo Fisher Scientific) coupled to an Exploris 480 Orbitrap mass spectrometer (Thermo Fisher Scientific). Full MS spectra were acquired at a resolution of 120,000, and the most abundant multiply charged precursor ions were selected for higher-energy collision dissociation (HCD) tandem mass spectrometry.

Raw data were processed using PEAKS X Pro against the UniProt *Bivalvia* database. Carbamidomethylation of cysteine residues was specified as a fixed modification, whereas N-terminal acetylation, methionine oxidation, asparagine/glutamine deamidation, and serine/threonine/tyrosine phosphorylation were considered variable modifications. Protein identifications were validated in ScaffoldDDA Viewer v6.7.4 using peptide- and protein-level false discovery rates of 1% and requiring at least two unique peptides. Proteins supported by a single spectral count or showing fewer than two additional spectral counts relative to the corresponding control were excluded. Fold change was calculated and then normalized with log base 2.

Common interactors identified in the different datasets were determined using BLAST+ (blastp) with a reciprocal best-hit strategy, retaining matches with ≥ 50% alignment coverage and ≥ 35% sequence identity. Functional annotation was performed using UniProt Gene Ontology (GO) terms and KEGG Orthology (KO) assignments generated with BlastKOALA (60). GO categories were summarized using WEGO 2.0 (61), and graphical analyses were performed in R (v4.5.2) using the ggplot2, dplyr, forcats, stringr, and scales packages (62).

### Structural and sequence analyses

The three-dimensional structures of *M. edulis* and *V. ellipsiformis* MORF proteins were predicted using AlphaFold 3 (63). Putative functional motifs, sorting signals, and subcellular localization were predicted with DeepLoc 2.1 (64), whereas intrinsically disordered regions were identified using flDPnn (65).

## RESULTS

### Histological characterization of reproductive and gametogenic stages in *M. edulis* and *V. ellipsiformis*

To establish the reproductive context for MORF localization and functional analyses, we first characterized male gonadal organization by hematoxylin and eosin staining. In *M. edulis*, males displayed fully mature gonads, with acini densely packed with mature spermatozoa. Distinct stages of spermatogenesis, including spermatogonia, spermatocytes, and spermatids, were also observed, indicating ongoing gamete production. Acrosomal structures were visible in spermatozoa when cells were not highly compacted (Fig. 1A-C).

**Figure 1.**
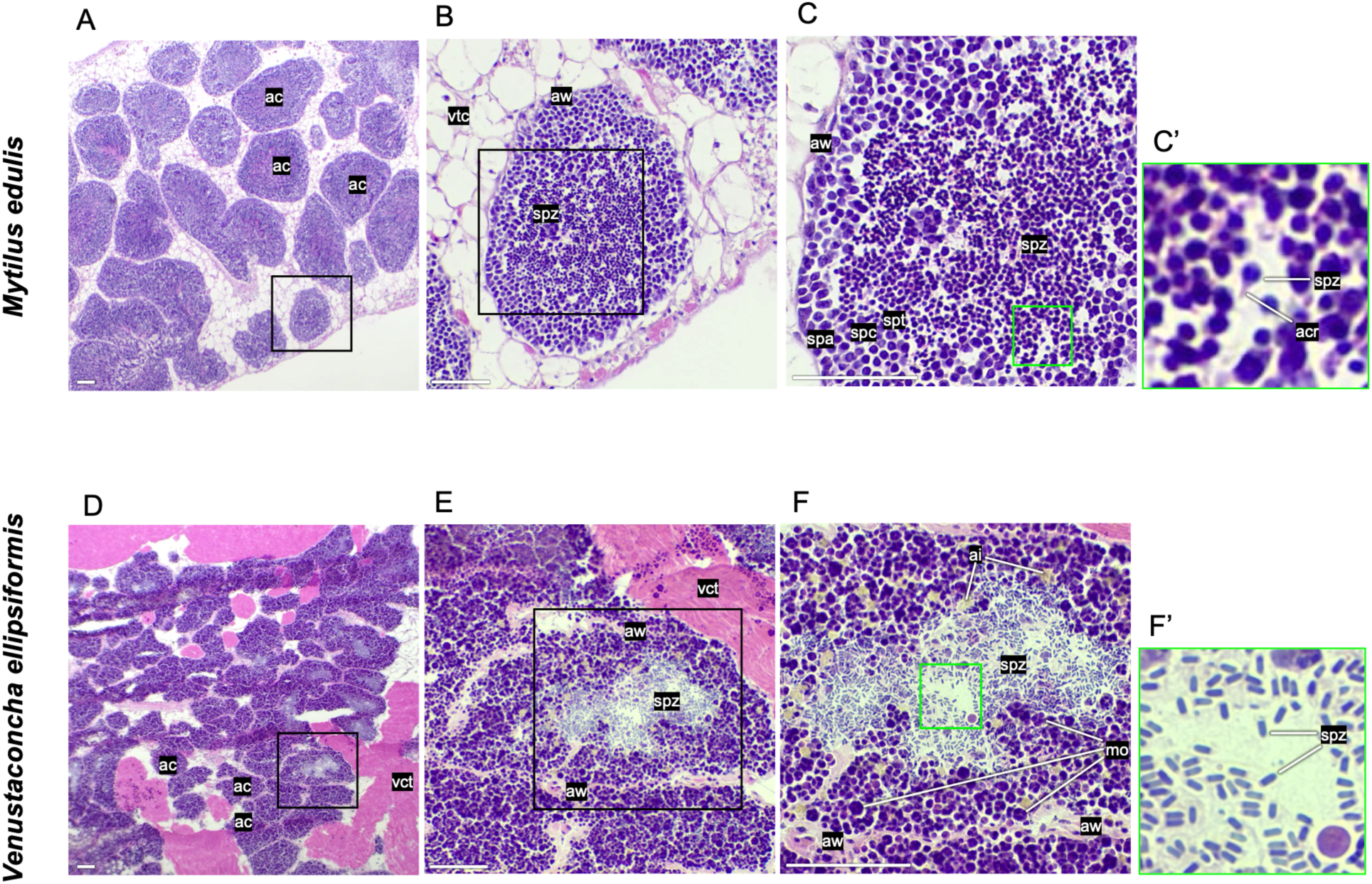
Hematoxylin and eosin (HE)-stained gonad sections from male *Mytilus edulis* and *Venustaconcha ellipsiformis.* Hematoxylin stains cell nuclei purple and eosin stains the extracellular matrix and the cytoplasm pink. (A-C) Gonad sections from a mature male *M. edulis.* (C’) Higher-magnification of the green box in (C). (D-F) Gonad sections from a mature male *V. ellipsiformis,* showing the atypical pathway of spermatogenesis. (F’) Higher-magnification of the green box in (F). Abbreviations: ac, acini; acr, acrosome; ai, amorphous inclusion; aw, acinus wall; mo, morulae; spa, spermatogonia; spc, spermatocystes; spt, spermatids; spz, spermatozoa; vct, visceral conjunctive tissue. Scale bars: 50 µm.

In *V. ellipsiformis*, males displayed variable reproductive states, ranging from recently spawned gonads with depleted follicles or tubules to ripe gonads containing mature spermatozoa. All four males analyzed displayed the atypical spermatogenic pathway previously described for this species, characterized by the presence of morulae-like structures at different development stages and amorphous inclusions (18). Mature spermatozoa were predominantly localized in the lumen of the acini (Fig. 1D-F).

### MORF localization in male gonads of *M. edulis* and *V. ellipsiformis*

We next examined MORF localization throughout spermatogenesis using immunohistochemistry combined with ATP5A staining as a mitochondrial marker. In *M. edulis*, MORF was detected across all stages of male germ cell development, from spermatogonia to mature spermatozoa, with spatial overlap with ATP5A signal (Fig. 2A-F; SI Appendix Fig. S1). This mitochondrial association was maintained throughout spermatocyte and spermatid differentiation and was particularly evident in mature spermatozoa. Notably, MORF was additionally detected within the fully formed acrosome, a region devoid of ATP5A staining (Fig. 2D-F).

**Figure 2.**
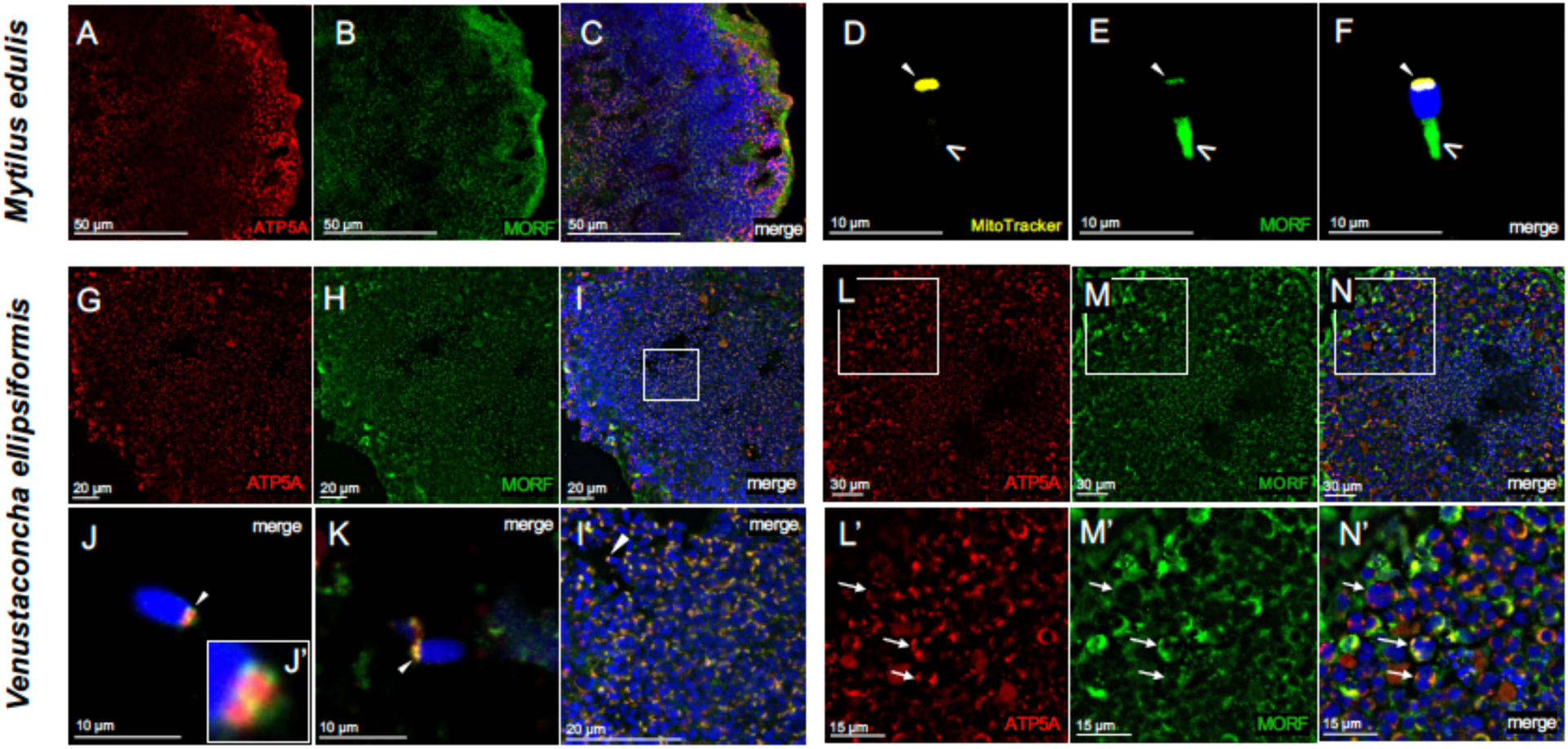
MORF subcellular localization in male germ cells and gonadal tissues of *Mytilus edulis* and *Venustaconcha ellipsiformis*. (A-C) *M. edulis* IHC. (D-F) Isolated mature sperm from *M. edulis* obtained from immunocytochemistry experiments described in (17). (G-N) *V. ellipsiformis* IHC. (I’) magnification of the white box in (I), (J’) magnification of the sperm mitochondrial rosette in (J), (L’) magnification of the white box in (L), (M’) magnification of the white box in (M) and (N’) magnification of the white box in (N). (J, K) Isolated mature sperm from *V. ellipsiformis* IHC. Anti-ATP5A shows mitochondria in red, anti-MORF shows MORF in green and DAPI displays nuclei in blue. White arrows point to morulae, white triangles to mitochondria and white v-shapes to the *M. edulis* sperm acrosome.

Occasional MORF immunoreactivity was observed within the acinar lumen (SI Appendix, Fig. S1). To determine whether this signal reflected secretion of MORF, conditioned media from cultured *M. edulis* male germ cells were analyzed by western blotting. MORF was detected in cellular extracts but was absent from culture media, like ATP5A (SI Appendix, Fig. S2), indicating that extracellular MORF signal was unlikely to result from active secretion under the tested conditions.

In *V. ellipsiformis*, MORF was detected in mature spermatozoa (but not in the acrosome) and colocalized with ATP5A in the mitochondrial region (Fig. 2G-K). MORF signal was also observed surrounding sperm mitochondria (Fig. 2J-K; SI Appendix, Movie S1). MORF and ATP5A signals were also detected in other acinar cell populations, including morula-like cells, although signal intensity varied among cells (Fig. 2L-N).

### MORF influences mitochondrial organization during spermatid differentiation

To investigate whether MORF contributes to mitochondrial organization during spermatid development, primary *M. edulis* male germ cell cultures were treated with anti-MORF antibodies, and mitochondrial organization was visualized using MitoTracker staining. Antibody-mediated neutralization of MORF altered the distribution of mitochondrial configuration in developing *M. edulis* spermatids (SI Appendix, Table S1). The estimated frequency of spermatids containing four rather than five mitochondria was 6.6% in the pre-immune serum control, 10.8% in the secondary-antibody control and 23.2% following anti-MORF treatment. Treatment had a significant effect on the probability of observing four-mitochondria spermatids (GLMM: χ^2^ = 23.42, df = 2, *P* = 8.21 × 10^-6^) (Fig. 3A). Relative to the pre-immune serum control, anti-MORF treatment increased the odds of observing four-mitochondria spermatids (odds ratio = 4.30, *P* = 1.26 × 10^-5^), whereas the secondary antibody-only control (Ab2) did not differ significantly from the pre-immune serum control (odds ratio = 1.71, *P* = 0.14). The enrichment of four-mitochondria spermatids following MORF neutralization was also supported by Fisher’s exact test on pooled counts (*P* = 1.23 × 10^-6^). Together, these results associate MORF neutralization with a shift away from the canonic five-mitochondria configuration during spermatid differentiation.

**Figure 3.**
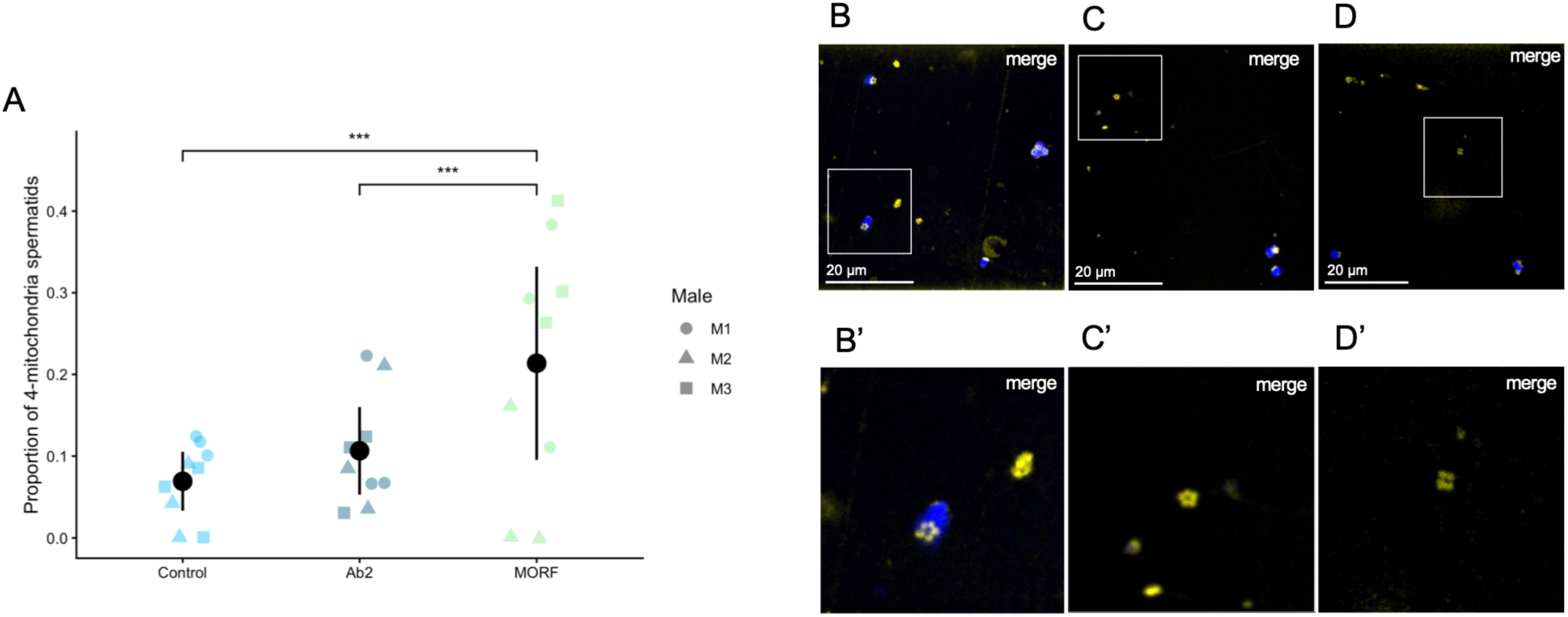
MORF association with mitochondrial rosette formation in cultured spermatid maturation in *M. edulis*. (A) Dot plot representation of the four-mitochondria spermatid proportion in each treatment (pre-immune control, secondary antibody-only control [Ab2] and anti-MORF) in three *M. edulis* males. *** *P* < 0.001. (B-D) Immunocytochemistry on cultured *M. edulis* male germ cells. (B’) magnification of the white box in (B), (C’) magnification of the white box in (C) and (D’) magnification of the white box in (D). (B-B’) 5-mitochondria spermatid. (C-C’) Free five-mitochondria rosette. (D-D’) Free four-mitochondria rosette. MitoTracker shows them in yellow, and DAPI displays nuclei in blue.

These findings were supported by the observation and quantification of free but metabolically active mitochondrial rosettes (Fig. 2B-D; SI Appendix, Table S1). Free mitochondrial rosettes containing four mitochondria were observed at higher frequency in anti-MORF-treated samples than in controls, whereas no comparable increase was detected for free rosettes containing five mitochondria, reinforcing the conclusion that MORF neutralization interferes with the formation of the typical five-mitochondria assemblies.

### Predicted structural features and intrinsic disorder of MORF proteins

To explore potential molecular determinants underlying MORF function, we compared predicted structural features of *M. edulis* and *V. ellipsiformis* MORF protein sequences (Fig. 4A) using AlphaFold3 and disorder prediction approaches. The two MORF proteins displayed markedly different predicted architectures. *V. ellipsiformis* MORF was predicted to adopt a structured α-helical conformation containing helix-turn-helix (HTH)- and leucine zipper (LZ)-like features, whereas *M. edulis* MORF showed overall low structural confidence (pLDDT < 50) and was predicted to contain predominantly antiparallel β-sheet regions (Fig. 4B).

**Figure 4.**
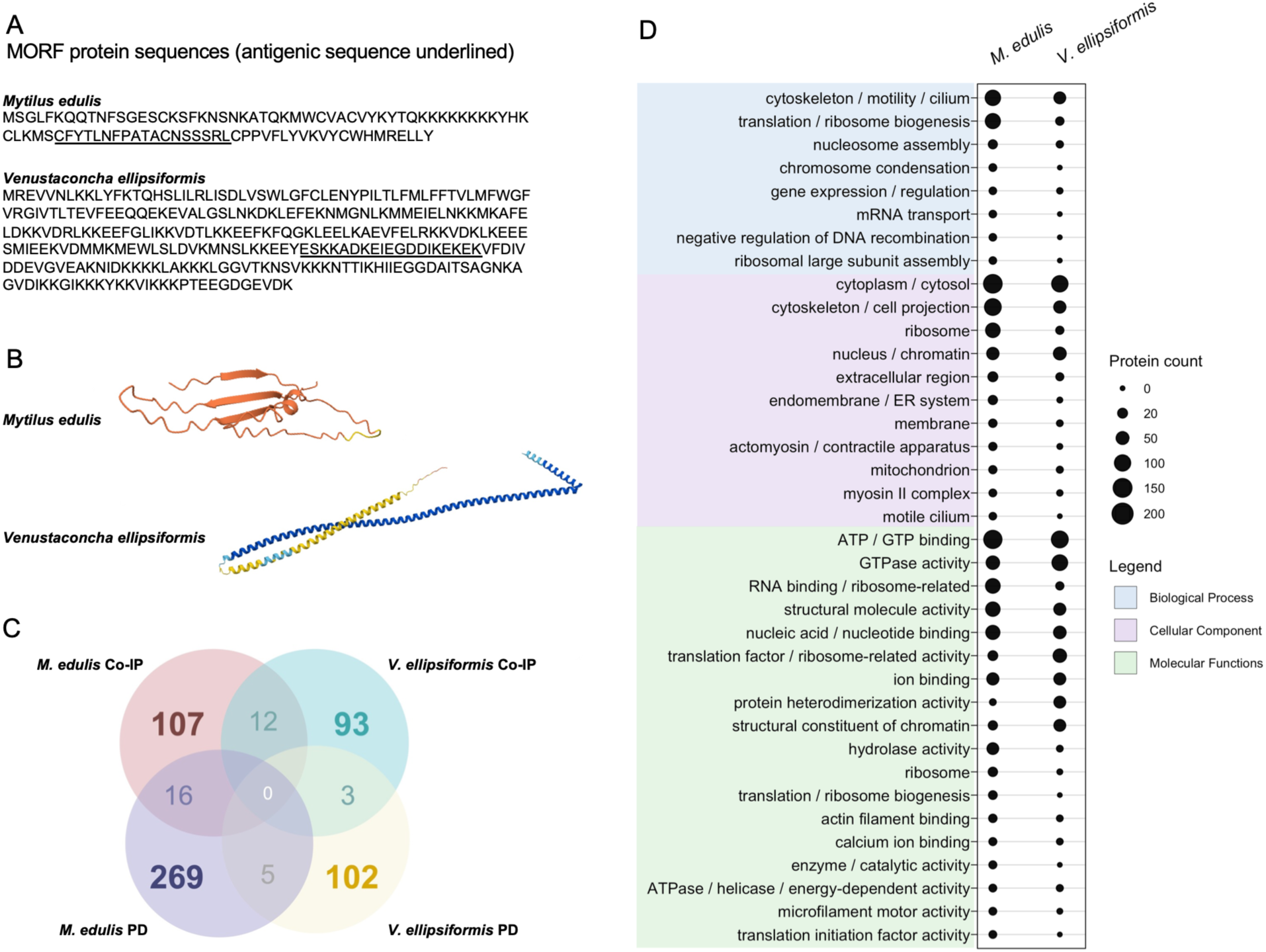
Analyses of MORF interacting partners and *in silico* analyses of MORF protein sequences from *Venustaconcha ellipsiformis* and *Mytilus edulis*. (A) Amino acid sequences of *V. ellipsiformis* and *M. edulis* MORF with antigenic peptide underlined. (B) Most confident 3D structure predictions generated by AlphaFold for *V. ellipsiformis* and *M. edulis* MORF. (C) Venn diagram illustrating the distribution of protein counts identified by co-immunoprecipitation (Co-IP) and GST pull-down assay (PD) in *M. edulis* and *V. ellipsiformis* samples. Values represent protein unique to each group of shared between groups. (D) Bubble plot representation of the most represented Gene Ontology (GO) terms for each GO category (i.e., Biological Process, Cellular Component of Molecular Function) among MORF-interacting proteins identified in *M. edulis* and *V. ellipsiformis*.

Subcellular localization predictions using DeepLoc 2.1 suggested species-specific differences in MORF topology. *V. ellipsiformis* MORF contained a predicted transmembrane segment with high confidence (probability = 0.906), with endoplasmic reticulum localization receiving the highest prediction score (0.857). In contrast, *M. edulis* MORF predictions were less conclusive, with similar probabilities for membrane-associated and soluble states and highest scores for nuclear and cytoplasmic localization (SI Appendix, Fig. S3 and Tables S2-S5).

Because both MORF sequences displayed features compatible with intrinsically disordered regions (IDRs), intrinsic disorder was further evaluated using flDPnn. In *M. edulis* MORF, two major IDRs were predicted: an N-terminal region encompassing the first 10 residues and an internal ∼20-amino-acid region containing a poly-lysine stretch of eight residues. These regions were associated with predicted nucleic acid- and protein-binding potential. In *V. ellipsiformis* MORF, two C-terminal IDRs of approximately 25 amino acids each were predicted, separated by a short ordered region. These regions were also associated with predicted nucleic acid- and protein-binding potential. Poly-lysine stretches were detected in both species, suggesting potential contributions of disordered, charged regions to MORF molecular interactions (SI Appendix, Fig. S4).

### Proteomic analyses identify MORF-associated candidates linked to extracellular matrix organization, cytoskeletal architecture and proteostasis

To define the molecular environment of MORF, we characterized MORF-associated protein complexes from male gonads of *M. edulis* and *V. ellipsiformis* using complementary co-immunoprecipitation (Co-IP) and GST pull-down (PD) approaches coupled with tandem mass spectrometry (SI Appendix, Dataset S1). The specificity of MORF enrichment (and biochemical confirmation of *in vivo* expression) was confirmed by the detection of multiple MORF-derived peptides in our co-immunoprecipitation samples, including four peptides corresponding to *V. ellipsiformis* MORF and three peptides corresponding to *M. edulis* MORF (SI Appendix, Fig. S5).

Across both species and approaches, 542 candidate MORF-associated proteins were identified, including 350 proteins in *M. edulis* and 192 proteins in *V. ellipsiformis*. Sixteen proteins were common between Co-IP and pull-down datasets in *M. edulis*, whereas three overlapped in *V. ellipsiformis* (Fig. 4C; SI Appendix, Tables S6 and S7). Twelve proteins were shared between species in Co-IP datasets and five in pull-down datasets, with no protein common to both species and both approaches (Fig. 4C; SI Appendix, Tables S8 and S9). Despite this limited overlap, the identification of high-confidence interactors using the peptide- and enrichment-weighted metric Δ(n) × FC^2^, where Δ(n) represents the treatment-to-control peptide ratio and FC^2^ the squared fold change, highlighted three main functional categories: extracellular matrix components (ECM), cytoskeletal architecture and motor activity, and regulation of translation and proteostasis (Tables 1,2).

**Table 1.** Top 10 detected proteins in mass spectrometry analyses obtained from *Mytilus edulis* co-immunoprecipitations and pull-down assays for two males, ranged by mean score.

| **Method** | **Protein Name** | **UniProt Accession Number** | **Male 1** |  |  |  |  | **Male 2** |  |  |  |  | **mean FC** | **mean log2 FC** | **mean score** |
|---|---|---|---|---|---|---|---|---|---|---|---|---|---|---|---|
|  |  |  | **IP PC** | **CTL PC** | **FC** | **log2 FC** | **score** | **IP PC** | **CTL PC** | **FC** | **log2 FC** | **score** |  |  |  |
| **Co-IP** | LAMC1 | A0A8S3T270 | 2 | 0 | 3 | 1.58 | 18 | 11 | 0 | 12 | 3.58 | 1584 | 7.5 | 2.58 | 801 |
|  | Laminin, gamma 1 | A0A8B6GQ75 | 2 | 0 | 3 | 1.58 | 18 | 11 | 0 | 12 | 3.58 | 1584 | 7.5 | 2.58 | 801 |
|  | Laminin, gamma 1 | A0A8B6GPC6 | 2 | 0 | 3 | 1.58 | 18 | 11 | 0 | 12 | 3.58 | 1584 | 7.5 | 2.58 | 801 |
|  | LAMC1 | A0A6J8A8F3 | 2 | 0 | 3 | 1.58 | 18 | 9 | 0 | 10 | 3.32 | 900 | 6.5 | 2.45 | 459 |
|  | SOD1 | A0A6J8DSI4 | 5 | 0 | 6 | 2.58 | 180 | 4 | 1 | 2.5 | 1.32 | 18.75 | 4.25 | 1.95 | 99.36 |
|  | 40S ribosomal protein S12 | A0A6J8EWE8 | 3 | 0 | 4 | 2 | 48 | 8 | 1 | 4.5 | 2.17 | 141.75 | 4.25 | 2.08 | 94.88 |
|  | Large subunit ribosomal protein | A0A8B6EMQ0 | 11 | 5 | 2 | 1 | 24 | 4 | 0 | 5 | 2.32 | 100 | 3.5 | 1.66 | 62 |
|  | Laminin, alpha 3/5 | A0A8B6G9Q8 | 3 | 0 | 4 | 2 | 48 | 6 | 2 | 2.33 | 1.22 | 21.78 | 3.17 | 1.61 | 34.89 |
|  | Large ribosomal subunit protein uL11m | A0A6J8BLS5 | 9 | 4 | 2 | 1 | 20 | 3 | 0 | 4 | 2 | 48 | 3 | 1.5 | 34 |
|  | ubiquitinyl hydrolase 1 | A0A210QZJ1 | 3 | 0 | 4 | 2 | 48 | 2 | 0 | 3 | 1.58 | 18 | 3.5 | 1.79 | 33 |
| **PD** | Collagen, type VI, alpha | A0A8B6H5N9 | 12 | 0 | 13 | 3.70 | 2,028 | 35 | 0 | 36 | 5.17 | 45,360 | 24.5 | 4.44 | 23,694 |
|  | MYH6_7 | A0A8S3SNY1 | 16 | 8 | 1.89 | 0.92 | 28.54 | 29 | 0 | 30 | 4.91 | 26,100 | 15.94 | 2.91 | 13,064.27 |
|  | Uncharacterized protein | A0A8B6EYF2 | 9 | 0 | 10 | 3.32 | 900 | 22 | 0 | 23 | 4.52 | 11,638 | 16.5 | 3.92 | 6,269 |
|  | DNA-directed RNA polymerase subunit beta' | A0A2T6N717 | 13 | 0 | 14 | 3.81 | 2,548 | 20 | 0 | 21 | 4.39 | 8,820 | 17.5 | 4.10 | 5,684 |
|  | Uncharacterized protein | A0A8S3RD63 | 10 | 0 | 11 | 3.46 | 1,210 | 20 | 0 | 21 | 4.39 | 8,820 | 16 | 3.93 | 5,015 |
|  | Uncharacterized protein | A0A8B6HIW0 | 10 | 7 | 1.38 | 0.46 | 5.67 | 19 | 0 | 20 | 4.32 | 7,600 | 10.69 | 2.39 | 3,802.83 |
|  | Core-binding (CB) domain-containing protein | A0A8B6G6U8 | 2 | 0 | 3 | 1.58 | 18 | 17 | 0 | 18 | 4.17 | 5,508 | 10.5 | 2.88 | 2,763 |
|  | Uncharacterized protein | A0A8S3REC1 | 2 | 0 | 3 | 1.58 | 18 | 17 | 0 | 18 | 4.17 | 5,508 | 10.5 | 2.88 | 2,763 |
|  | Paramyosin,Myosin-7 | A0A8S3RM96 | 16 | 2 | 5.67 | 2.50 | 449.56 | 16 | 0 | 17 | 4.09 | 4,624 | 11.33 | 3.29 | 2,536.78 |
|  | Elongation factor Tu | A0AAE0VMK0 | 13 | 0 | 14 | 3.81 | 2,548 | 11 | 0 | 12 | 3.58 | 1,584 | 13 | 3.70 | 2,066 |

Abbreviations: Co-IP: co-immunoprecipitation; FC: fold change; log2FC: log base 2-fold change; IP: immunoprecipitation; CTL: control; PC: peptide count.

**Table 2.**
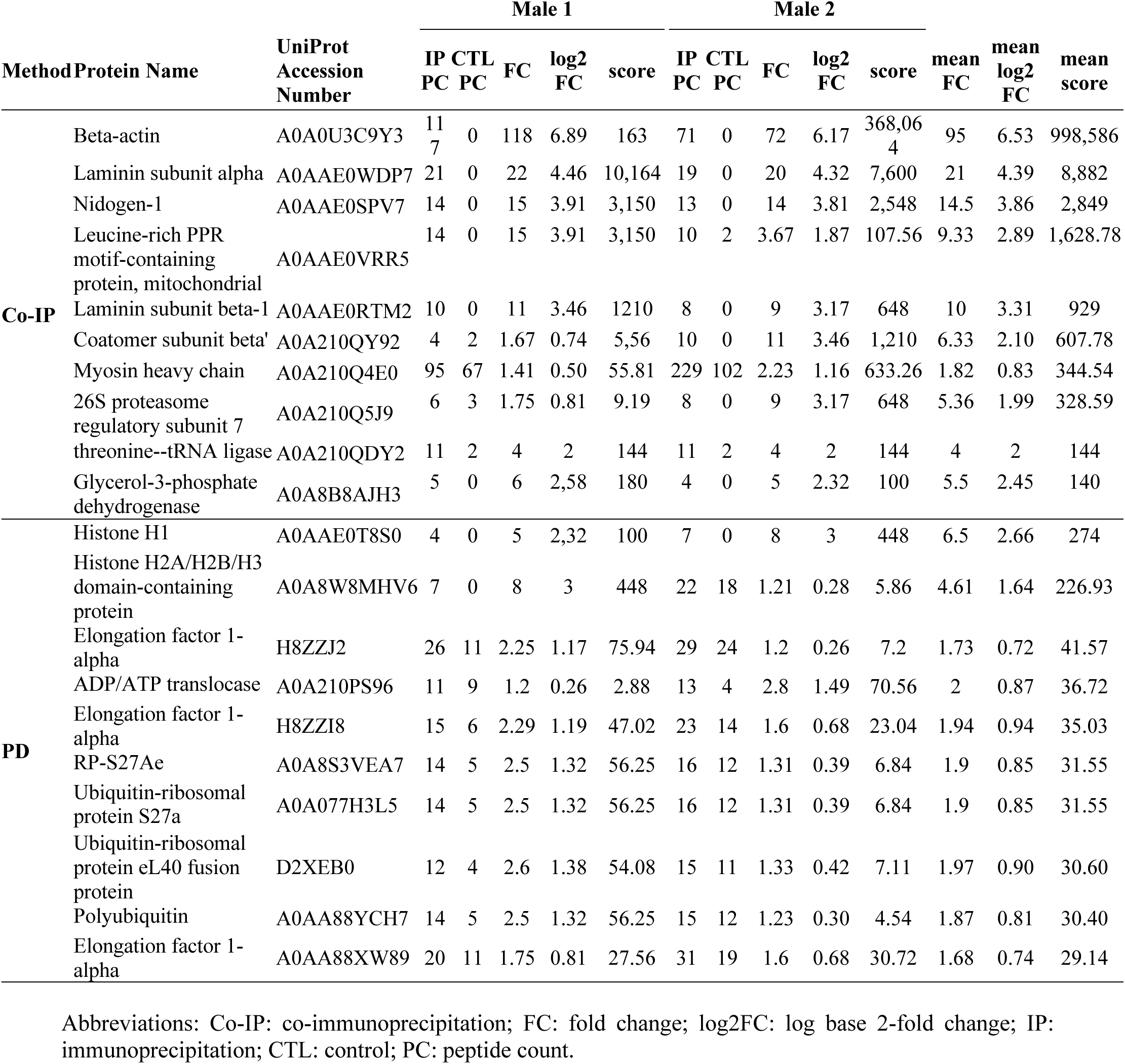
Top 10 detected proteins in mass spectrometry analyses obtained from *Venustaconcha ellipsiformis* co-immunoprecipitations and pull-down assays for two males, ranged by mean score.

| Method | Protein Name | UniProt Accession Number | Male 1 |  |  |  |  | Male 2 |  |  |  |  |  |  |  |
|---|---|---|---|---|---|---|---|---|---|---|---|---|---|---|---|
|  |  |  | **IP PC** | **CTL PC** | **FC** | **log2 FC** | **score** | **IP PC** | **CTL PC** | **FC** | **log2 FC** | **score** | **mean FC** | **mean log2 FC** | **mean score** |
| **Co-IP** | Beta-actin | A0A0U3C9Y3 | 117 | 0 | 118 | 6.89 | 163 | 71 | 0 | 72 | 6.17 | 368,064 | 95 | 6.53 | 998,586 |
|  | Laminin subunit alpha | A0AAE0WDP7 | 21 | 0 | 22 | 4.46 | 10,164 | 19 | 0 | 20 | 4.32 | 7,600 | 21 | 4.39 | 8,882 |
|  | Nidogen-1 | A0AAE0SPV7 | 14 | 0 | 15 | 3.91 | 3,150 | 13 | 0 | 14 | 3.81 | 2,548 | 14.5 | 3.86 | 2,849 |
|  | Leucine-rich PPR motif-containing protein, mitochondrial | A0AAE0VRR5 | 14 | 0 | 15 | 3.91 | 3,150 | 10 | 2 | 3.67 | 1.87 | 107.56 | 9.33 | 2.89 | 1,628.78 |
|  | Laminin subunit beta-1 | A0AAE0RTM2 | 10 | 0 | 11 | 3.46 | 1210 | 8 | 0 | 9 | 3.17 | 648 | 10 | 3.31 | 929 |
|  | Coatomer subunit beta' | A0A210QY92 | 4 | 2 | 1.67 | 0.74 | 5,56 | 10 | 0 | 11 | 3.46 | 1,210 | 6.33 | 2.10 | 607.78 |
|  | Myosin heavy chain | A0A210Q4E0 | 95 | 67 | 1.41 | 0.50 | 55.81 | 229 | 102 | 2.23 | 1.16 | 633.26 | 1.82 | 0.83 | 344.54 |
|  | 26S proteasome regulatory subunit 7 | A0A210Q5J9 | 6 | 3 | 1.75 | 0.81 | 9.19 | 8 | 0 | 9 | 3.17 | 648 | 5.36 | 1.99 | 328.59 |
|  | threonine--tRNA ligase | A0A210QDY2 | 11 | 2 | 4 | 2 | 144 | 11 | 2 | 4 | 2 | 144 | 4 | 2 | 144 |
|  | Glycerol-3-phosphate dehydrogenase | A0A8B8AJH3 | 5 | 0 | 6 | 2,58 | 180 | 4 | 0 | 5 | 2.32 | 100 | 5.5 | 2.45 | 140 |
| **PD** | Histone H1 | A0AAE0T8S0 | 4 | 0 | 5 | 2,32 | 100 | 7 | 0 | 8 | 3 | 448 | 6.5 | 2.66 | 274 |
|  | Histone H2A/H2B/H3 domain-containing protein | A0A8W8MHV6 | 7 | 0 | 8 | 3 | 448 | 22 | 18 | 1.21 | 0.28 | 5.86 | 4.61 | 1.64 | 226.93 |
|  | Elongation factor 1-alpha | H8ZZJ2 | 26 | 11 | 2.25 | 1.17 | 75.94 | 29 | 24 | 1.2 | 0.26 | 7.2 | 1.73 | 0.72 | 41.57 |
|  | ADP/ATP translocase | A0A210PS96 | 11 | 9 | 1.2 | 0.26 | 2.88 | 13 | 4 | 2.8 | 1.49 | 70.56 | 2 | 0.87 | 36.72 |
|  | Elongation factor 1-alpha | H8ZZI8 | 15 | 6 | 2.29 | 1.19 | 47.02 | 23 | 14 | 1.6 | 0.68 | 23.04 | 1.94 | 0.94 | 35.03 |
|  | RP-S27Ae | A0A8S3VEA7 | 14 | 5 | 2.5 | 1.32 | 56.25 | 16 | 12 | 1.31 | 0.39 | 6.84 | 1.9 | 0.85 | 31.55 |
|  | Ubiquitin-ribosomal protein S27a | A0A077H3L5 | 14 | 5 | 2.5 | 1.32 | 56.25 | 16 | 12 | 1.31 | 0.39 | 6.84 | 1.9 | 0.85 | 31.55 |
|  | Ubiquitin-ribosomal protein eL40 fusion protein | D2XEB0 | 12 | 4 | 2.6 | 1.38 | 54.08 | 15 | 11 | 1.33 | 0.42 | 7.11 | 1.97 | 0.90 | 30.60 |
|  | Polyubiquitin | A0AA88YCH7 | 14 | 5 | 2.5 | 1.32 | 56.25 | 15 | 12 | 1.23 | 0.30 | 4.54 | 1.87 | 0.81 | 30.40 |
|  | Elongation factor 1-alpha | A0AA88XW89 | 20 | 11 | 1.75 | 0.81 | 27.56 | 31 | 19 | 1.6 | 0.68 | 30.72 | 1.68 | 0.74 | 29.14 |

Abbreviations: Co-IP: co-immunoprecipitation; FC: fold change; log2FC: log base 2-fold change; IP: immunoprecipitation; CTL: control; PC: peptide count.

Extracellular matrix proteins were prominent among the MORF-associated candidates identified by proteomic analyses. In *M. edulis*, several laminin subunits ranked among the most enriched proteins in the co-IP dataset, whereas the *V. ellipsiformis* dataset included laminin subunits, nidogen and cartilage matrix proteins. Collagen VI was also identified as a prominent candidate in the *M. edulis* pull-down dataset (Tables 1,2). Importantly, the association between MORF and laminin was independently supported by western blot analyses of gonad homogenates and GST-MORF pull-down assays in both species (SI Appendix, Fig. S6 and S7). Immunolocalization further revealed laminin at the basal membrane of *M. edulis* male gonadal (SI Appendix, Fig. S8). These complementary results support the reproducibility of the MORF-laminin association, while the remaining proteins identified by proteomics should be considered candidate interactors pending independent validation.

Gene Ontology enrichment analysis revealed consistent functional signatures across species and experimental approaches (Fig. 4D; Tables 1,2; SI Appendix, Dataset S1). Identified proteins were primarily associated with genetic information processing, including translation and RNA metabolism, as well as cytoskeletal organization, motor activity, and macromolecular complex assembly. Molecular function categories were enriched for structural components and nucleic acid-binding activities, whereas ECM-associated proteins, particularly laminins, nidogens, and collagens, were recurrently detected, especially in Co-IP datasets.

KEGG Orthology analysis confirmed enrichment for genetic information processing as the dominant category across *M. edulis* Co-IP/PD and *V. ellipsiformis* PD datasets, followed by environmental information processing, cellular processes, and organismal systems (SI Appendix, Fig. S9). Together, these findings identify MORF as a conserved component of a broad molecular network connecting mitochondrial organization with cytoskeletal remodeling, extracellular matrix-associated structures, and cellular homeostasis during spermatogenesis.

## DISCUSSION

The principal functional finding of this study is that antibody-mediated neutralization of MORF is associated with a disruption of the canonical mitochondrial organization of developing *M. edulis* spermatids. Previous studies established the male-specific expression of MORF and its association with sperm mitochondria (16, 17) but did not directly test its contribution to mitochondrial organization during spermatogenesis. The increased frequency of four-mitochondria configurations following MORF neutralization therefore extends these descriptive observations by providing experimental evidence for a functional contribution of MORF to sperm development. Although the the molecular basis of this phenotype remains unresolved, our findings implicate MORF in the formation or maintenance of the characteristic mitochondrial arrangement in sperm.

Beyond advancing our understanding of MORF function, these findings reinforce the emerging view that mitochondrial ORFan genes represent biologically meaningful components of cellular systems rather than mere evolutionary curiosities. They further support the hypothesis that lineage-specific mitochondrial genes can evolve novel cellular roles, contributing to the diversification and evolution of organismal traits, including those associated with reproductive systems.

### MORF is associated with sperm mitochondrial organization throughout spermatogenesis

Histological analyses confirmed the marked differences in spermatogenic organization between the two species examined. In *M. edulis*, spermatogenesis followed the canonical pattern described for most metazoans, with successive germ cell stages arranged along the acini and mature spermatozoa accumulating within the lumen (19). In contrast, all *V. ellipsiformis* males displayed the atypical spermatogenic pathway previously described for unionids, characterized by morula-like germ cell aggregates and amorphous inclusions interpreted as degenerating morulae (18). Because specimens were collected during autumn, this organization likely reflects the seasonal reproductive cycle of unionids and has been proposed to represent an energetically reduced mode of spermatogenesis during overwintering (18).

Despite these architectural differences, MORF exhibited a conserved localization pattern. In both species, the protein was detected continuously from the earliest germ cells through mature spermatozoa, indicating that its expression is maintained throughout spermatogenesis rather than being restricted to terminal sperm differentiation. In *M. edulis*, these observations extend previous reports that localized MORF to late spermatids and mature sperm (16, 17), and demonstrate that its mitochondrial association persists throughout germ cell development. Similarly, MORF was detected in the atypical germ cell aggregates of *V. ellipsiformis*, suggesting that its recruitment is conserved despite profound differences in spermatogenic architecture.

One of the most significant findings of this study is that neutralization of MORF altered the stereotypical organization of sperm mitochondria during spermatid formation. Blocking MORF significantly increased the frequency of spermatids and free mitochondrial rosettes containing four mitochondria instead of the canonical five-mitochondria arrangement characteristic of *M. edulis*. The four-mitochondria configurations retained detectable MitoTracker Red CMXRos staining, indicating that the observed phenotype was not accompanied by a complete loss of mitochondrial membrane potential under the conditions examined. This observation is consistent with an alteration of mitochondrial organization rather than whole mitochondrial depolarization, although additional assays would be required to determine whether other aspects of mitochondrial function were affected. To our knowledge, this represents the first functional evidence that a mitochondrial ORFan contributes to mitochondrial architecture during gametogenesis.

This interpretation is further supported by MORF localization. In both species, MORF was associated with the mitochondrial region of developing and mature sperm cells, although its immunofluorescence signal did not uniformly overlap with ATP5A. This partial overlap may reflect differences in the subcellular distribution of the two proteins, including the association of MORF with specific mitochondrial subregions or adjacent structures. In *V. ellipsiformis*, MORF immunoreactivity was also observed surrounding sperm mitochondria (16, present study), raising the possibility that MORF is enriched at particular mitochondrial surfaces or at interfaces between neighboring mitochondria. However, conventional immunofluorescence does not resolve the precise molecular topology of MORF, and the available data do not establish its direct localization at mitochondria-mitochondria contact sites. Higher-resolution imaging will be required to determine the precise localization of MORF relative to mitochondrial membranes and neighboring organelles.

Mitochondrial clustering during spermatogenesis requires coordinated regulation of mitochondrial dynamics, organelle positioning, and cytoskeletal transport (20, 21), and disruption of these processes compromises sperm development and male fertility in humans (22). The functional effect of MORF neutralization therefore provides experimental support for previous hypotheses that MORF participates in mitochondrial organization during sperm maturation (16).

In mature *M. edulis* spermatozoa, MORF was additionally detected within the acrosome, confirming previous observations (16). Although the acrosomal signal was less clearly resolved in *V. ellipsiformis*, likely owing to the acrosome’s tight apposition to the nucleus and minimal subacrosomal material head, combined with methodological resolution limits, mitochondrial localization remained unequivocal in both species. The acrosomal localization may indicate an additional role in acrosome biogenesis or fertilization-related processes, although it may also reflect secondary recruitment of mitochondrial proteins during sperm differentiation. Functional analyses focused specifically on acrosome formation will be required to distinguish between these possibilities.

An additional observation was the occasional detection of MORF immunoreactivity outside germ cells, particularly along the acinar periphery. Western blot analyses of conditioned media did not support active secretion of MORF under the culture conditions tested, arguing against constitutive extracellular release. Nevertheless, the in vitro culture system may not fully reproduce the physiological environment of the gonad. Alternatively, the observed staining could represent antibody retention within densely packed tissues, as previously described in incompletely permeabilized specimens (23). A third possibility is that MORF associates with relatively insoluble extracellular matrix components rather than being freely secreted. This hypothesis is particularly intriguing given the strong enrichment of laminins, collagens, and nidogens among MORF-associated proteins identified in the present study (see below), and the striking similarity between MORF immunostaining and reported localization patterns of extracellular matrix proteins within gonadal tissues (24, 25). Although further work will be required to distinguish among these possibilities, the convergence of localization and interactome data suggests that MORF may participate in structural interfaces linking germ cells to their extracellular environment.

### Conserved regulatory features despite structural divergence

Although AlphaFold3 predicted markedly different structures for the MORF proteins of *M. edulis* and *V. ellipsiformis*, several lines of evidence suggest that these proteins may share a common functional architecture. The *V. ellipsiformis* model exhibited a well-defined α-helical fold containing predicted helix-turn-helix (HTH) and leucine zipper-like (LZ) motifs, features frequently associated with nucleic acid binding and transcriptional regulation (26, 27). In contrast, the *M. edulis* model showed globally low prediction confidence (pLDDT < 50) and lacked a stable three-dimensional fold. While this could reflect genuine structural flexibility, it is also consistent with the well-recognized difficulty of predicting ORFan protein structures because of their absence of detectable homologs and atypical evolutionary histories (28–30). Consequently, the apparent structural divergence between species should be interpreted cautiously until experimental structural data become available.

Despite these differences in predicted folds, our analyses point to several conserved features that are likely to be functionally more informative. Both MORF proteins contain prominent intrinsically disordered regions (IDRs) enriched in predicted nucleic acid- and protein-binding residues. IDRs are increasingly recognized as central components of regulatory proteins because they enable conformational plasticity, multivalent interactions, and dynamic assembly of molecular complexes (31, 32). Such proteins typically function as interaction hubs rather than rigid structural components, allowing them to participate in multiple cellular processes in a context-dependent manner. This interpretation is remarkably consistent with the extensive MORF interactome identified in the present study.

The presence of poly-lysine motifs embedded within these disordered regions further supports this hypothesis. Polybasic sequences promote interactions with negatively charged molecules, including nucleic acids, biological membranes, and acidic protein surfaces, and contribute to targeting proteins to subcellular compartments such as the nucleus, endoplasmic reticulum, and mitochondria (33–36). Rather than representing isolated sequence motifs, these lysine-rich regions may therefore provide the molecular basis for the diverse interaction profile observed experimentally.

Subcellular localization predictions are also consistent with a multifunctional regulatory role. Although DeepLoc assigned different intracellular localizations to the two MORF proteins, neither prediction contradicts the experimental observations. *V. ellipsiformis* MORF displayed stronger association with membrane compartments, particularly the endoplasmic reticulum, whereas *M. edulis* MORF showed higher probabilities for cytoplasmic and nuclear localization. These predictions reflect sequence features associated with subcellular localization rather than the site of protein synthesis. As such, their agreement with the experimentally observed mitochondrial localization lies in supporting the potential for context-dependent localization rather than defining a single primary intracellular compartment. Such apparent versatility is characteristic of intrinsically disordered proteins, whose functions and localization are often dictated by interaction partners and cellular context rather than constitutive targeting signals (37). Accordingly, these computational predictions should be viewed as complementary to, rather than replacing, the experimental evidence demonstrating mitochondrial localization throughout spermatogenesis.

Collectively, these observations suggest that MORF is best viewed not as a conventional structural protein but as a dynamic molecular scaffold capable of coordinating multiple interaction networks during germ cell differentiation. Importantly, the ability of MORF to interact with extramitochondrial proteins does not necessarily imply that the protein is freely distributed throughout the cell. Instead, MORF may act as a mitochondria-associated platform that transiently engages cytoskeletal, extracellular matrix-associated, and regulatory proteins at sites where mitochondrial positioning and cellular remodeling are coordinated. Such a function would be consistent with the properties of intrinsically disordered proteins, which frequently mediate context-dependent and multivalent interactions through flexible regions rather than through a single stable structural domain. In this framework, MORF could serve as a molecular interface linking mitochondria to broader cellular networks involved in organelle positioning, cell architecture, and germ cell differentiation. Such functional plasticity would be consistent with the experimental demonstration that MORF neutralization perturbs mitochondrial organization without abolishing mitochondrial activity, implying a role in the spatial organization of molecular complexes rather than in core mitochondrial metabolism itself.

### MORF is a dynamic interaction hub protein coordinating mitochondrial organization during spermatogenesis

The MORF interactome provides a mechanistic framework for interpreting its role during spermatogenesis. Across both species, co-immunoprecipitation and GST pull-down experiments identified hundreds of candidate MORF-associated proteins spanning diverse cellular functions. Although the identities of MORF-associated candidates showed limited overlap across species and experimental approaches, the datasets shared several broad functional annotations including cytoskeletal organization, extracellular matrix-associated processes and protein homeostasis. This convergence suggests that MORF may be associated with similar cellular processes in both species, despite extensive sequence divergence. However, the limited overlap at the level of individual proteins prevents us from concluding that the underlying molecular interaction networks are conserved. Further experiments will be required to establish whether these shared functional annotations reflect conserved underlying mechanisms or distinct species-specific interactions.

The larger number of proteins recovered by GST pull-down compared with co-immunoprecipitation is itself informative. Whereas co-immunoprecipitation preferentially captures stable protein complexes, recombinant bait proteins used in pull-down assays also recover transient and moderate-affinity interactions (38–40). Together with the predicted intrinsically disordered regions, this interaction profile is consistent with MORF behaving as a hub protein, capable of forming dynamic, multivalent interaction networks instead of stable stoichiometric complexes (41–43). Such behavior is characteristic of regulatory proteins that coordinate multiple cellular processes through transient molecular interactions.

Three functional modules emerged consistently across species and experimental approaches: proteins involved in genetic information processing, cytoskeletal organization, and extracellular matrix (ECM) components.

The first module comprised proteins involved in translation, RNA metabolism, ribosome biogenesis, and protein homeostasis. Spermatogenesis is accompanied by profound transcriptional and translational reprogramming that requires tight temporal coordination between mitochondrial activity and gene expression (44, 45). Rather than suggesting that MORF directly regulates transcription or translation, these interactions likely reflect its integration within the molecular environment supporting germ cell differentiation. In this context, MORF may contribute to coordinating mitochondrial activity with the biosynthetic demands associated with sperm maturation.

The second module comprised cytoskeletal proteins, including myosins, paramyosins, tropomyosins, and other motor-associated components. This finding is particularly compelling in light of the functional neutralization experiments. During spermatogenesis, mytilid and unionid sperm mitochondria undergo extensive spatial reorganization to form a distinct five-mitochondria rosette in the sperm midpiece (18, 19). In diverse organisms, mitochondrial positioning depends on coordinated interactions between cytoskeletal networks, molecular motors, and mitochondrial tethering factors (20, 21). Our demonstration that MORF neutralization selectively increased the frequency of “rarer”, four-mitochondria assemblies, while preserving mitochondrial activity (as shown by MitoTracker), strongly supports a role in the spatial organization of mitochondria. The localization of MORF surrounding sperm mitochondria, together with its interaction with cytoskeletal proteins, therefore suggests that MORF contributes to the assembly or stabilization of the mitochondrial rosette by coordinating organelle positioning during spermatid maturation. These observations provide functional support for previous bioinformatic predictions implicating MORF in cytoskeletal organization (15, 46), and complement recent transcriptomic evidence linking cytoskeleton-associated genes with sex-specific mitochondrial inheritance in DUI embryos (47).

The third module, and perhaps the most unexpected, involved extracellular matrix proteins. Laminins, collagens, and nidogens consistently ranked among the highest-confidence MORF interactors in both species, and the MORF-laminin association was independently validated by western blotting. ECM proteins are core basement membrane components providing structural scaffolding and regulating adhesion and differentiation (48, 49). In bivalves, germ cells develop upon basement membranes enriched in laminins, collagens, and nidogens (18, 50–54). Beyond their structural role, laminins have recently been implicated in promoting mitochondrial clustering and mitochondria-endoplasmic reticulum contacts in other systems (55). Although it remains speculative whether a similar mechanism operates during bivalve spermatogenesis, the convergence of ECM proteins, cytoskeletal components, and mitochondrial localization within the MORF interactome suggests that these processes may be functionally linked. Because MORF localization was consistently associated with sperm mitochondria, the detection of ECM components among its interactors should not be interpreted as evidence that MORF functions as an extracellular protein. Rather, these interactions may reflect the existence of molecular bridges connecting mitochondria with adhesion and cytoskeletal networks within developing germ cells and their surrounding microenvironment.

Taken together, our data support a model in which MORF acts as a dynamic molecular scaffold positioned at the interface between mitochondria, cytoskeletal networks, and cell-matrix-associated signaling pathways. Rather than functioning as a structural component of mitochondria themselves, MORF may facilitate the coordinated assembly of molecular complexes required for mitochondrial positioning during spermatid differentiation. Such a role readily explains its intrinsically disordered architecture, broad interaction profile, and the selective disruption of mitochondrial organization observed following MORF neutralization.

More broadly, our findings substantially expand the functional repertoire currently attributed to mitochondrial ORFan genes. MORF is not simply a lineage-specific mitochondrial protein expressed during spermatogenesis; it is a functional regulator whose perturbation alters a defining cellular event of sperm differentiation. Because faithful organization of paternal mitochondria is central to doubly uniparental inheritance, the functional association between MORF and mitochondrial organization raises the possibility that this lineage-specific protein contributes to aspects of male reproductive biology relevant to paternal mitochondrial transmission. However, our experiments do not directly test mitochondrial inheritance across fertilization or embryogenesis. Determining whether MORF influences the retention or transmission of paternal mitochondria will require functional analyses extending beyond spermatid differentiation.

An intriguing parallel can be drawn with prohibitin, a conserved mitochondrial scaffold that regulates mitochondrial architecture, nucleoid organization, and spermatid differentiation while interacting with cytoskeletal and adhesion proteins (56, 57). Interestingly, a prohibitin-like protein emerged as a potential MORF interactor in *M. edulis*, representing an exciting direction for future investigation.

## CONCLUSION

Our results identify MORF as a mitochondrial-associated protein expressed throughout male germ cell development in two distantly related bivalves with doubly uniparental inheritance. In *M. edulis*, antibody-mediated neutralization of MORF alters the canonical mitochondrial organization of developing spermatids, providing functional evidence that this lineage-specific protein contributes to sperm development. Complementary proteomic analyses identify MORF-associated candidates linked to extracellular matrix organization, cytoskeletal architecture and protein homeostasis, and the association between MORF and laminin is supported by independent biochemical validation in both species. Although the precise molecular mechanisms remain unresolved, these findings connect a rapidly evolving mitochondrial protein to the organization of a specialized reproductive cell. More broadly, this study provides evidence that lineage-specific mitochondrial orphan genes can contribute to cellular processes beyond mitochondrial bioenergetics.

