## Supplementary Material for "A mitochondrial orphan protein contributes to sperm development in bivalves"

**FIGURES**

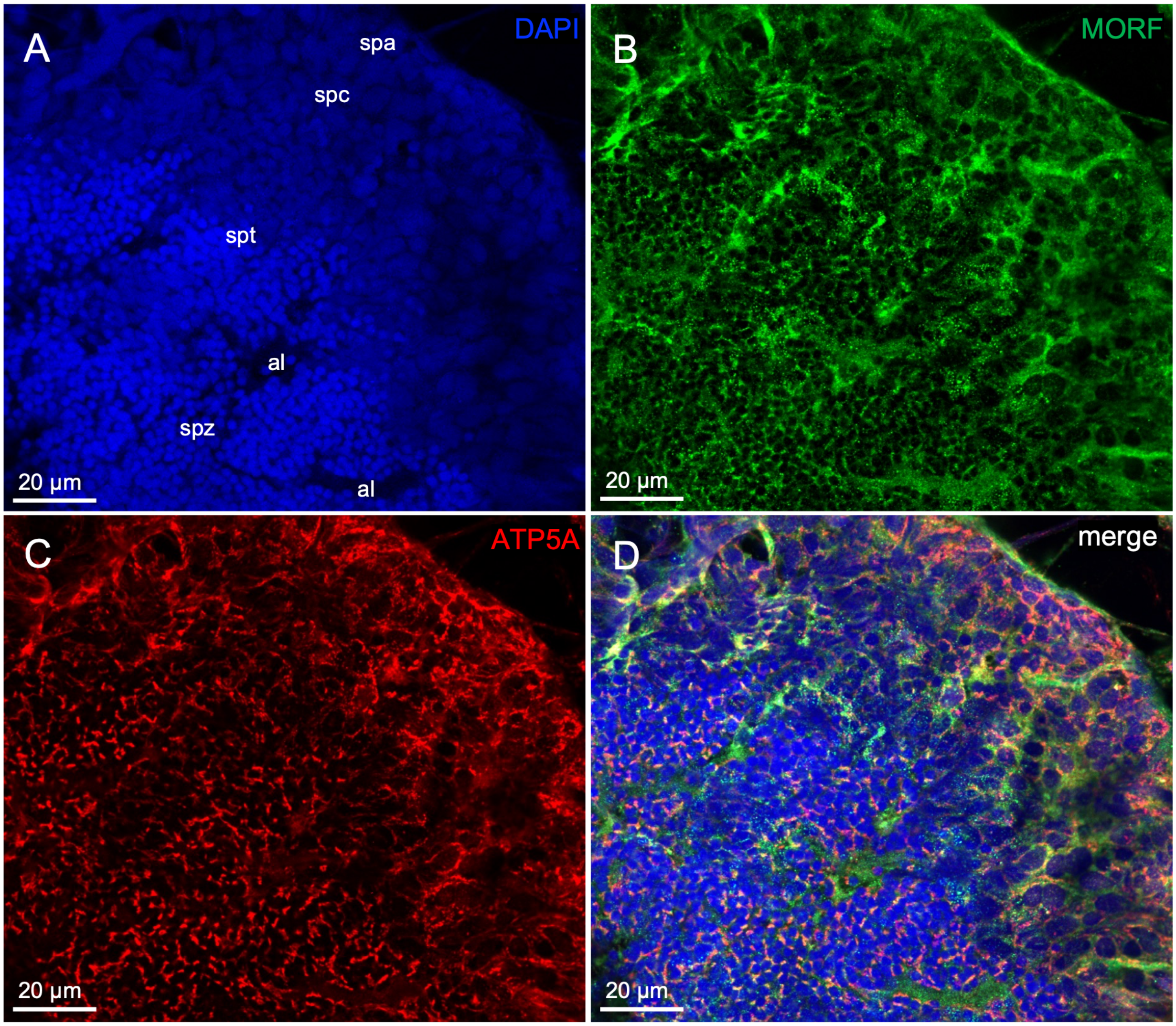


**Fig. S1. MORF localization revealed by immunohistochemistry in male gonadal tissues of *Mytilus edulis*.** (A) DAPI staining (blue) highlights nuclei within the acinus. (B) MORF immunofluorescence signal (green). (C) ATP5A immunofluorescence signal (red). (D) Merged image of the three channels. Abbreviations: al: acinar lumen; spa: spermatogonia; spc: spermatocyst; spt: spermatid; spz: spermatozoa.

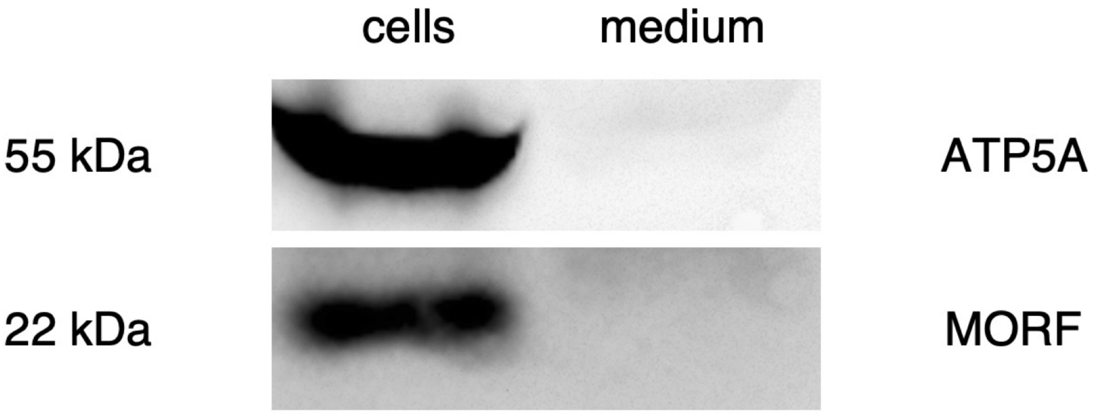


**Fig. S2. Western blot of *Mytilus edulis* male germ cell culture homogenates and culture medium using anti-MORF and anti-ATP5A antibodies.** ATP5A is detected at 55 kDa in *M. edulis* male gonad homogenates, and MORF at 22 kDa.

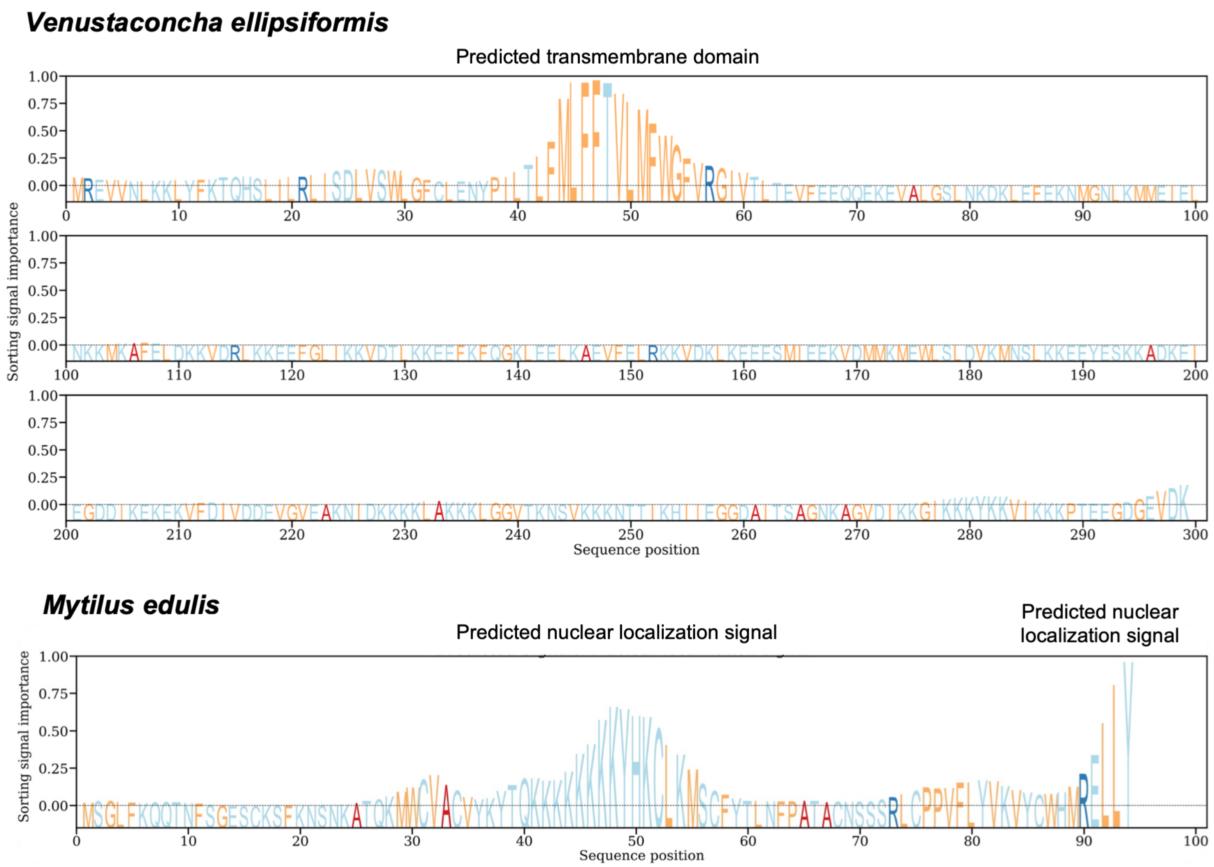


**Fig. S3. Predictions of potentially important functional regions using the DeepLoc 2.1 algorithm.**

### *Venustaconcha ellipsiformis*

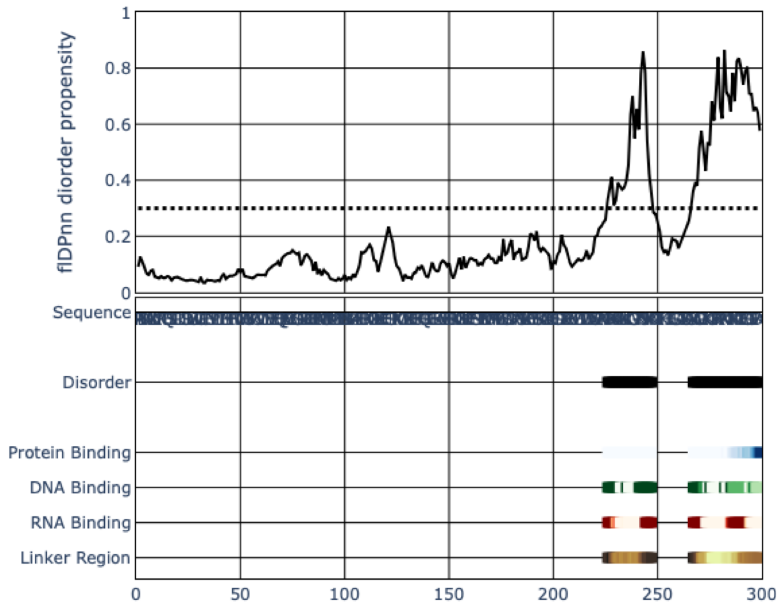


### *Mytilus edulis*

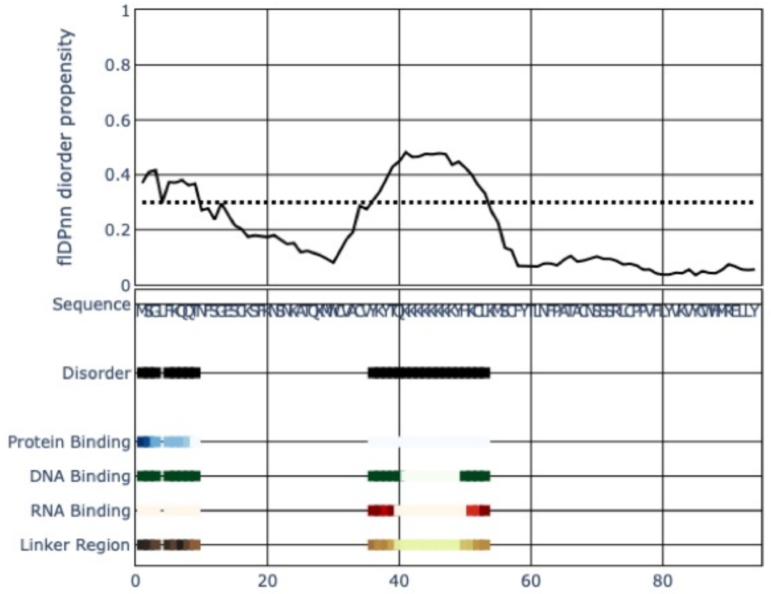


**Fig. S4. Prediction of intrinsically disordered regions (IDRs) in both MORF protein sequences using the flDPnn webserver.** Predicted functional roles are indicated for each disordered region.

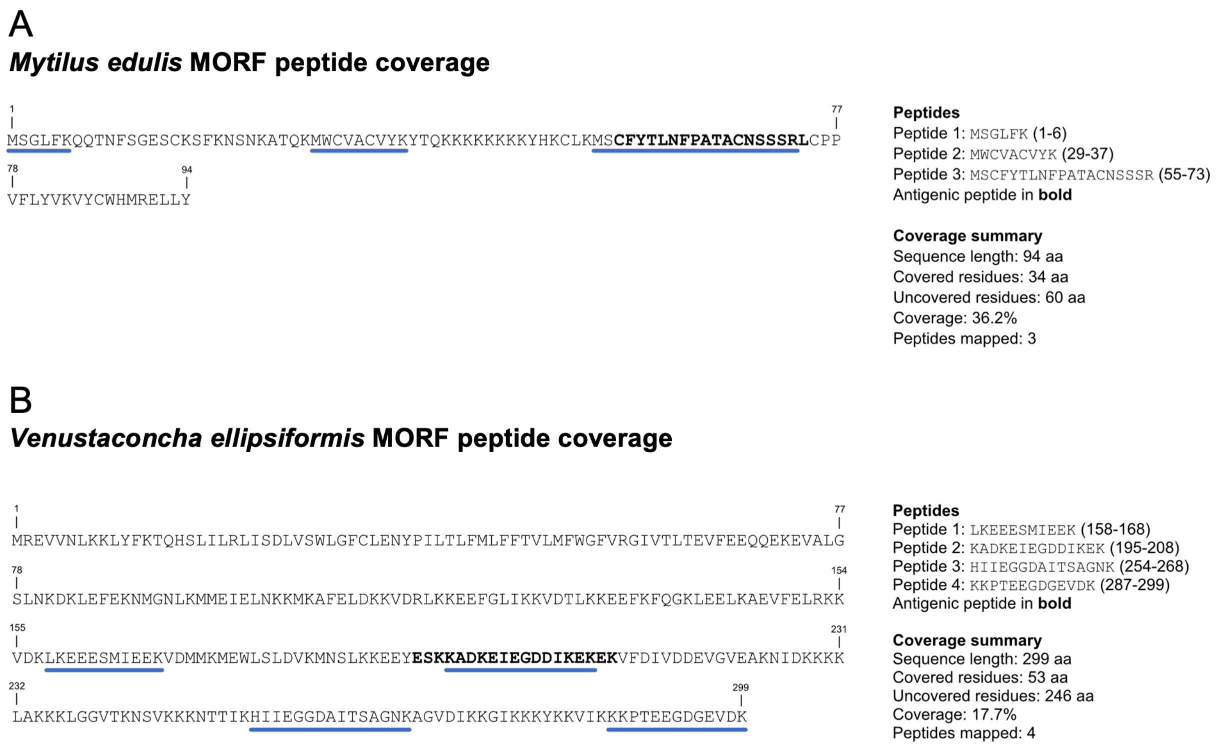


**Fig. S5. Unique peptides detected by tandem mass spectrometry for *Venustaconcha ellipsiformis* MORF and *Mytilus edulis* MORF.** (A) For *M. edulis* MORF, the three peptides were detected in a single sample. (B) For *V. ellipsiformis*, the peptide HIIEGGDAITSAGNK was detected in three different biological samples, and KADKEIEGDDIKEK and KKPTEEGDGEVDK in two, and finally LKEEESMIEEK in one.

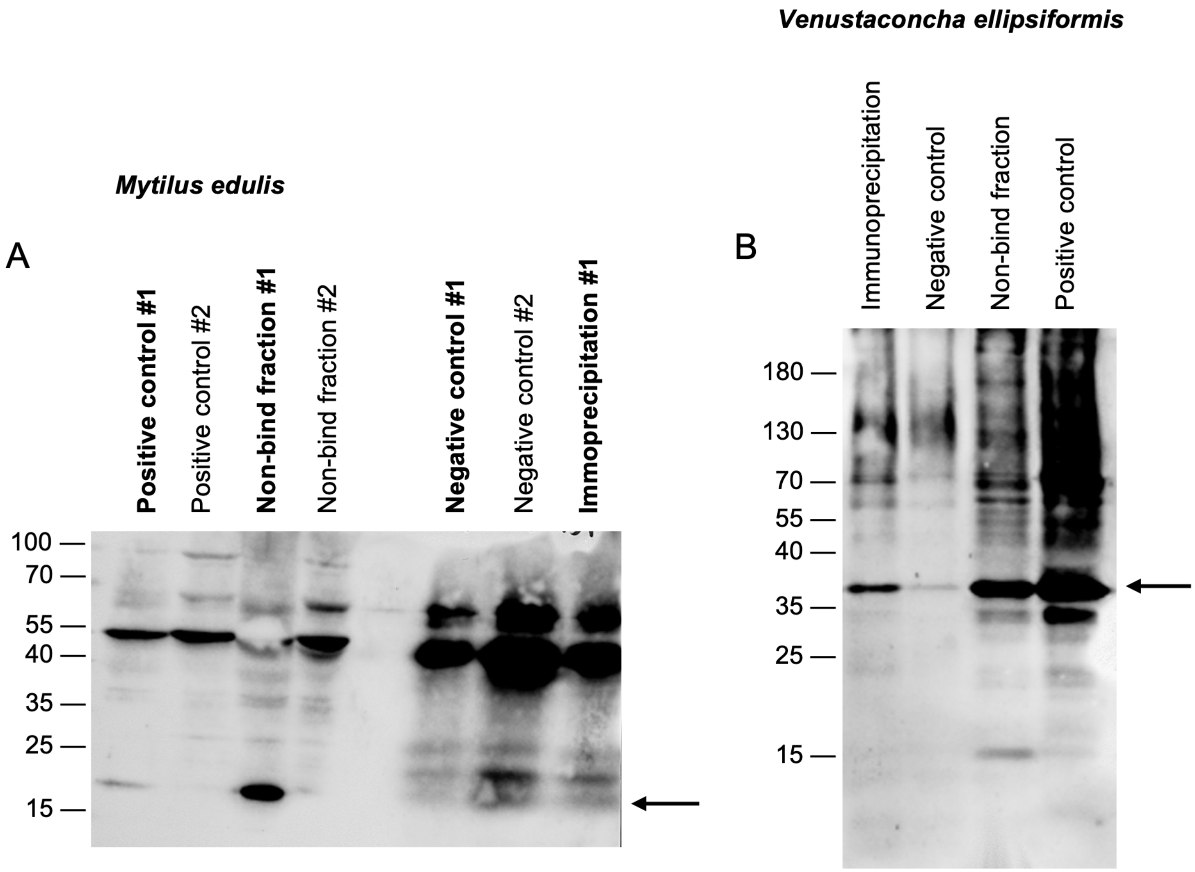


**Fig. S6. Complete membranes of western blotting confirming the immunoprecipitation of MORF.** (A) Western blotting of MORF from two *Mytilus edulis* males (#1 and #2). In #1, the arrow show that MORF is detected at 22 kDa in the positive control, the non-bind fraction. The band is more intense in the immunoprecipitation (IP) fraction than in the negative control, showing a successful IP. Higher smeared bands represent the heavy (proxy 50 kDa) and the light (proxy 25 kDa) chains. (B) Western blotting of MORF from *Venustaconcha ellipsiformis*. MORF is present at 36 kDa in all fractions, but at a lower concentration in the negative control, thus demonstrating a successful IP. The non-bind fraction represents the supernatant collected after the immunoprecipitation step and indicates the proportion of MORF (compared to the positive control) that did not bind the beads. Protein ladders are in kDa.

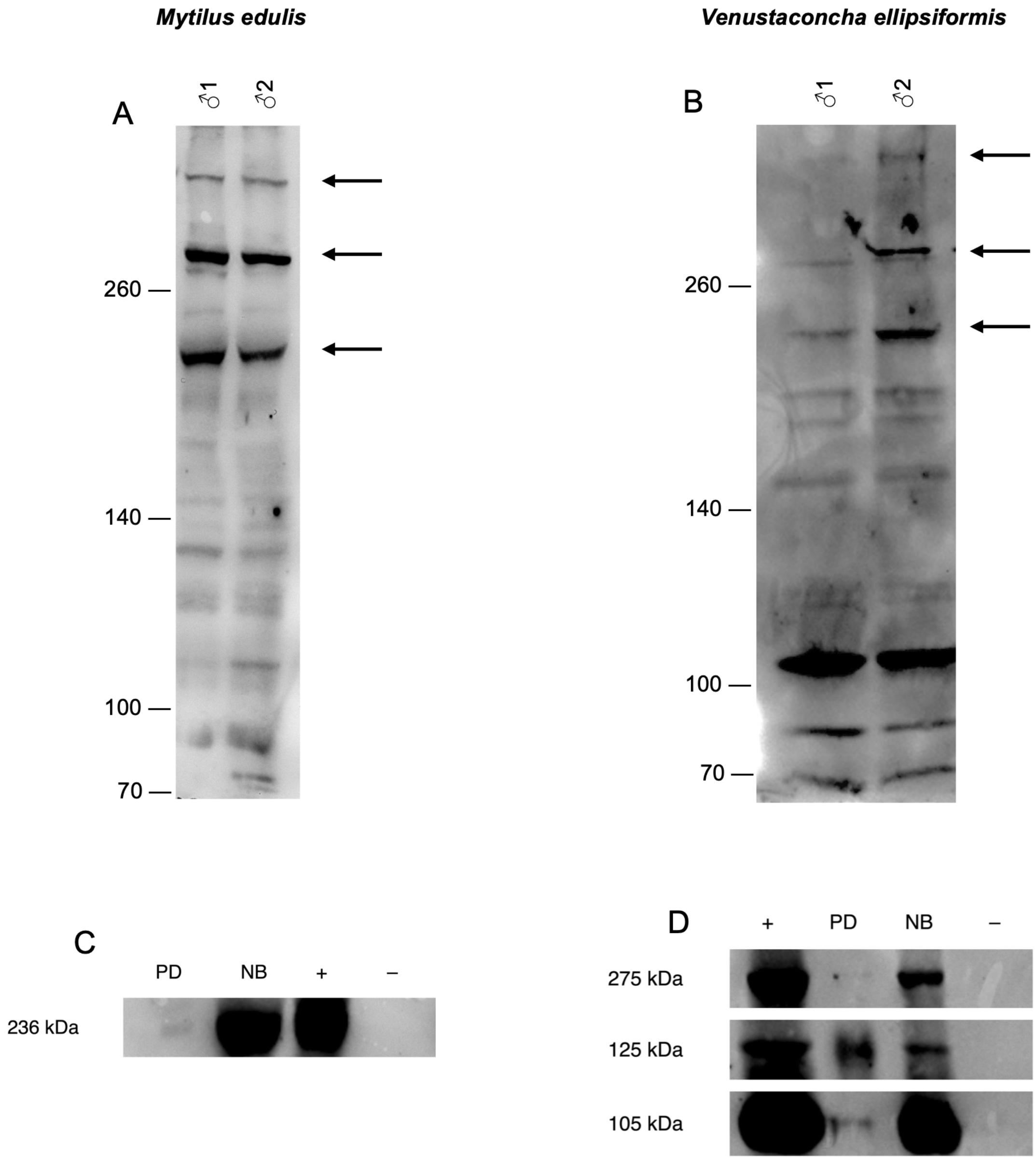


**Fig. S7. Western blotting of laminin from gonad homogenates and pull-down fraction from male *Mytilus edulis* and *Venustaconcha ellipsiformis* individuals.** (A) Western blotting of laminin in *Mytilus edulis* male gonad homogenates. (B) Western blotting of laminin from *Venustaconcha ellipsiformis* male gonad homogenates. For (A-B), three different bands (indicated by black arrows) were observed in two biological replicates in both species: from high to low kDa, alpha subunit (~305 kDa), beta subunit (~275 kDa) and gamma subunit (~236 kDa). (C) Western blotting of laminin following GST-MORF pull-down assays of male *Mytilus edulis* male gonad homogenates. A unique band was observed at proxy 236 kDa in all the fractions but the negative control. (D) Western blotting of laminin following GST-MORF pull-down assays of male *Venustaconcha ellipsiformis* male gonad homogenates. Three different bands were observed in all fractions but the negative control: at proxy 275 kDa, 125 kDa and 105 kDa. Abbreviations: PD: pull-down fraction; NB: non-bound fraction; +: positive control (lysate); –: negative control (GST alone). Protein ladders are in kDa.

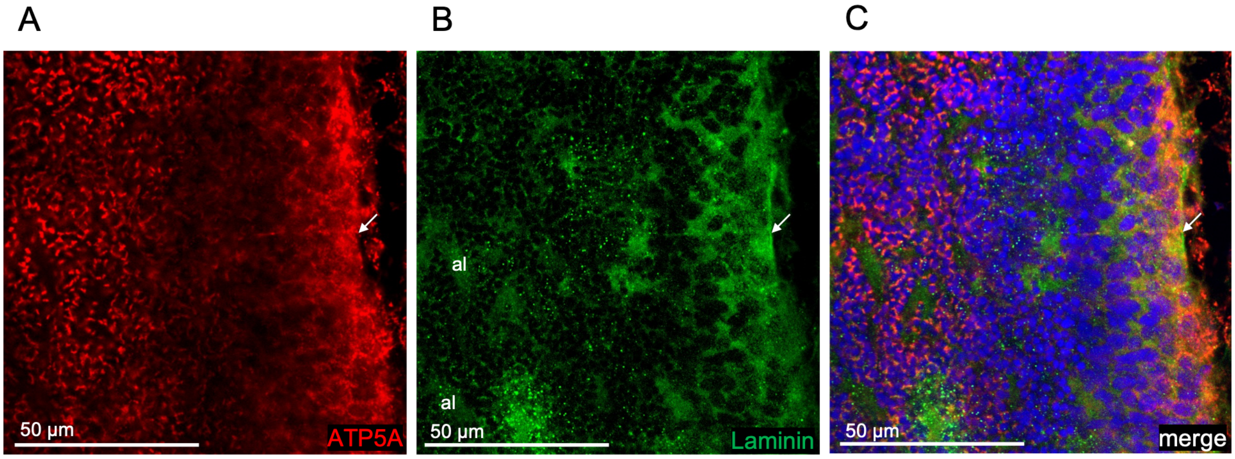


**Fig. S8. Immunohistochemical localization of laminin in male gonad sections of *Mytilus edulis*.** (A) Anti-ATP5A shows mitochondria in red. (B) Anti-laminin shows laminin in green. (C) Merge of red and green with DAPI displaying nuclei in blue. White arrows point to basal membranes of the acinus. Abbreviations: al: acinar lumen.

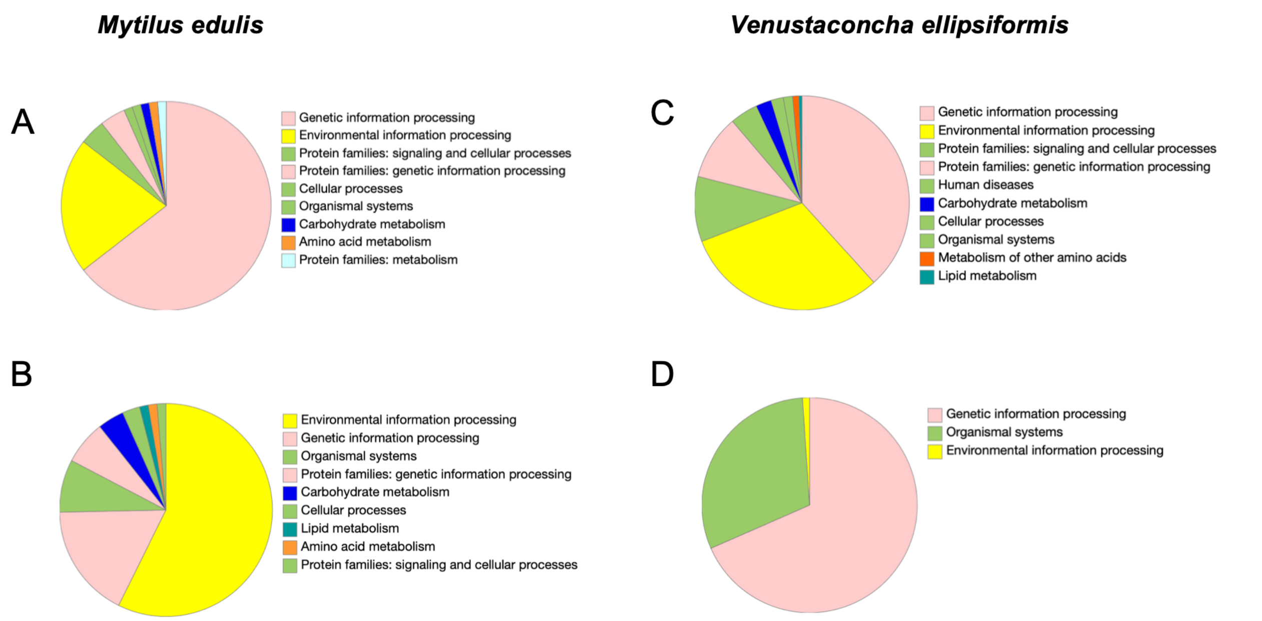


**Fig. S9. Pie chart representations of KEGG pathway annotations for MORF-interacting proteins identified by co-immunoprecipitations (Co-IP) and pull-down (PD) assays.** (A) *M. edulis* Co-IP; (B) *M. edulis* PD; (C) *V. ellipsiformis* Co-IP; (D) *V. ellipsiformis* PD.

**TABLES**

**Table S1. Distribution of spermatids according to mitochondrial content in three *Mytilus edulis* male germ cell cultures.**

| Male | Condition | Replicate | 5 mitochondria | 4 mitochondria | Unscorable spermatids | 4 free-mitochondria | 5 free-mitochondria |
|---|---|---|---|---|---|---|---|
| 1 | Control | A | 14 | 2 | 76 | 1 | 2 |
| | | B | 18 | 2 | 79 | 3 | 1 |
| | | C | 15 | 2 | 75 | 3 | 5 |
| | Ab2 | A | 14 | 1 | 51 | 1 | 3 |
| | | B | 14 | 4 | 96 | 1 | 4 |
| | | C | 14 | 1 | 64 | 2 | 2 |
| | MORF | A | 8 | 1 | 75 | 9 | 3 |
| | | B | 17 | 7 | 42 | 5 | 2 |
| | | C | 16 | 10 | 95 | 9 | 2 |
| 2 | Control | A | 20 | 2 | 50 | 1 | 3 |
| | | B | 23 | 1 | 53 | 0 | 5 |
| | | C | 14 | 0 | 53 | 0 | 3 |
| | Ab2 | A | 32 | 3 | 69 | 0 | 3 |
| | | B | 26 | 7 | 51 | 0 | 4 |
| | | C | 26 | 1 | 42 | 0 | 3 |
| | MORF | A | 22 | 0 | 64 | 3 | 10 |
| | | B | 16 | 0 | 68 | 2 | 4 |
| | | C | 21 | 4 | 57 | 0 | 4 |
| 3 | Control | A | 15 | 1 | 61 | 1 | 1 |
| | | B | 22 | 0 | 76 | 1 | 1 |
| | | C | 32 | 3 | 78 | 0 | 0 |
| | Ab2 | A | 30 | 1 | 88 | 0 | 2 |
| | | B | 24 | 3 | 106 | 0 | 1 |
| | | C | 21 | 3 | 124 | 0 | 3 |
| | MORF | A | 14 | 6 | 61 | 0 | 2 |
| | | B | 14 | 5 | 56 | 3 | 1 |
| | | C | 20 | 14 | 47 | 1 | 2 |

A, B and C are technical replicates; Control condition corresponds to the addition of pre-immun serum; Ab2 condition to the addition of a secondary antibody and MORF condition to the addition of anti-MORF antibody. Unscorable spermatids correspond to cells for which mitochondria number could not be determined because of cell orientation or nearby artifacts

**Table S2. Probabilities of *V. ellipsiformis* MORF localization calculated by Deeploc 2.1.** Bold probabilities are above the threshold.

| Localization | Cytoplasm | Nucleus | Extracellular | Cell membrane | Mitochondrion | Plastid | Endoplasmic reticulum | Lysosome / Vacuole | Golgi apparatus | Peroxisome |
|---|---|---|---|---|---|---|---|---|---|---|
| Probability | 0.2120 | 0.4244 | 0.2032 | 0.3206 | 0.1693 | 0.0362 | **0.8579** | 0.2146 | 0.3283 | 0.0659 |
| Probability threshold | 0.4761 | 0.5014 | 0.6173 | 0.5646 | 0.6220 | 0.6395 | 0.6090 | 0.5848 | 0.6494 | 0.7364 |

**Table S3**. **Probabilities of *V. ellipsiformis* MORF membrane association calculated by Deeploc 2.1.** Bold probabilities are above the threshold.

| Membrane association | Peripherical | Transmembrane | Lipid anchor | Soluble |
|---|---|---|---|---|
| Probability | 0.2640 | **0.9020** | 0.3080 | 0.3300 |
| Probability threshold | 0.6000 | 0.5100 | 0.8200 | 0.5000 |

**Table S4. Probabilities of *M. edulis* MORF localization calculated by Deeploc 2.1.** Bold probabilities are above the threshold.

| **Localization** | **Cytoplasm** | **Nucleus** | **Extracellular** | **Cell membrane** | **Mitochondrion** | **Plastid** | **Endoplasmic reticulum** | **Lysosome / Vacuole** | **Golgi apparatus** | **Peroxisome** |
|---|---|---|---|---|---|---|---|---|---|---|
| Probability | 0.4973 | **0.6032** | 0.3473 | 0.0480 | 0.3458 | 0.0021 | 0.2063 | 0.0481 | 0.0515 | 0.0018 |
| Probability threshold | 0.4761 | 0.5014 | 0.6173 | 0.5646 | 0.6220 | 0.6395 | 0.6090 | 0.5848 | 0.6494 | 0.7364 |

**Table S5. Probabilities of *M. edulis* MORF membranes association calculated by Deeploc 2.1.** Bold probabilities are above the threshold.

| Membrane association | Peripherical | Transmembrane | Lipid anchor | Soluble |
|---|---|---|---|---|
| Probability | 0.1510 | **0.5930** | 0.1000 | **0.6850** |
| Probability threshold | 0.6000 | 0.5100 | 0.8200 | 0.5000 |

**Table S6. UniProt accession numbers and associated protein names for common proteins found in *Mytilus edulis* co-immunoprecipitation and pull-down datasets.**

| *Mytilus edulis* Co-IP | | *Mytilus edulis* PD | |
|---|---|---|---|
| **Uniprot Accession Number** | **Protein names** | **Uniprot Accession Number** | **Protein name** |
| A0A077H3J9 | Small ribosomal protein uS9 | A0A077H3J9 | Small ribosomal protein uS9 |
| A0A077GZK0 | Small ribosomal protein uS5 | A0AAE0RRK1 | Small ribosomal protein uS5 |
| A0A8B6G1L2 | Uncharacterized protein | A0A8B6G1L2 | Uncharacterized protein |
| A0A077H0N7 | 60S ribosomal protein L7a | A0A077H0N7 | 60S ribosomal protein L7a |
| A0A6J8BHP0 | 40S ribosomal protein S4 | A0A6J8BHP0 | 40S ribosomal protein S4 |
| A0A8B6FY64 | Large ribosomal subunit protein uL15/eL18 domain-containg protein | A0A6J8AT14 | RP-L18e |
| D6BJL2 | Elongation factor 1-alpha | D6BJL2 | Elongation factor 1-alpha |
| A0A8S3SMY1 | Tubulin beta chain | A0A8S3SMY1 | Tubulin beta chain |
| A0A8B6EQG1 | Tubulin beta chain | A0A8B6EQG1 | Tubulin beta chain |
| A0A6J8CLH2 | Tubulin beta chain | A0A6J8CLH2 | Tubulin beta chain |
| A0A0K0YB09 | Transgelin | A0A0K0YB09 | Transgelin |
| A0A8B6FN82 | Tubulin beta chain | A0A8S3QXJ8 | Tubulin beta chain |
| A0A6J8AT33 | Nucleoside diphosphatase kinase 5 | A0A210QX70 | Nucleoside diphosphatase kinase 5 |
| A0A194ANJ0 | *(deleted)* Putative ribosomal protein L26 | A0A194ANJ0 | *(deleted)* Putative ribosomal protein L26 |

| | | | |
|---|---|---|---|
| A0A210QYL1 | Actin, cytoplasmic | A0A8S3R9S3 | Actin |
| A0A3G4ZJK8 | Histone H2B | A0A8B8D6D5 | Histone H2B |

Abbreviations: Co-IP: co-immunoprecipitation; PD: pull-down.

**Table S7. UniProt accession numbers and associated protein names for common proteins found in *Venustaconcha ellipsiformis* co-immunoprecipitation and pull-down datasets.**

| *Venustaconcha ellipsiformis* Co-IP | | *Venustaconcha ellipsiformis* PD | |
|---|---|---|---|
| **Uniprot Accession Number** | **Protein names** | **Uniprot Accession Number** | **Protein name** |
| I1W686 | *(deleted)* Histone H3 | A0A516V0F7 | *(deleted)* Histone H3 (fragment) |
| B3FEB6 | Histone H3 | V5L149 | *(deleted)* Histone H3 (fragment) |
| A0A5J6DQS7 | Heat shock protein 70 | A0A210QV24 | Endoplasmic reticulum chaperone BIP |

Abbreviations: Co-IP: co-immunoprecipitation; PD: pull-down.

**Table S8. UniProt accession numbers and associated protein names for common proteins found in *Mytilus edulis* and *Venustaconcha ellipsiformis* co-immunoprecipitation datasets.**

| *Mytilus edulis* Co-IP | | *Venustaconcha ellipsiformis* Co-IP | |
|---|---|---|---|
| **Uniprot Accession Number** | **Protein names** | **Uniprot Accession Number** | **Protein name** |
| A0AAE0WDP7 | Laminin subunit alpha-1 | A0A8B6G9Q8 | Laminin, alpha 3/5 |
| A0A8B6EEW8 | Tubulin beta chain | A0A8S3QXJ8 | Tubulin beta chain |
| A0A210PNZ3 | vesicule-fusing ATPase | A0A210PNZ3 | vesicule-fusing ATPase |
| A0A210QIC5 | Microtubule-associated protein futsch | A0A8B6FAG3 | Microtubule associated protein 1 |
| A0A8S3SMY1 | Tubulin beta chain | A0A8S3SMY1 | Tubulin beta chain |
| A0A8B6EQG1 | Tubulin beta chain | A0A8B6EQG1 | Tubulin beta chain |
| A0A6J8CLH2 | Tubulin beta chain | A0A6J8CLH2 | Tubulin beta chain |
| A0AA88YQH4 | Tubulin beta chain | A0AA88YQH4 | Tubulin beta chain |
| A0A8B8CKX3 | 60S ribosomal protein L3-like | A0A077GY47 | Ribosomal protein L3 |
| A0AAE0SGC7 | Actin | A0A8S3R9S3 | Actin |
| A0AAE0S1J9 | RRM domain-containing protein | A0A6J8ESL8 | HNRNPABD |
| A0A8B6GV24 | Tubulin/FtsZ 2-layer sandwich domain containing protein | A0A8B6GV24 | Tubulin/FtsZ 2-layer sandwich domain containing protein |

Abbreviations: Co-IP: co-immunoprecipitation.

**Table S9. UniProt accession numbers and associated protein names for common proteins found in *Mytilus edulis* and *Venustaconcha ellipsiformis* pull-down datasets.**

| *Mytilus edulis* PD | | *Venustaconcha ellipsiformis* PD | |
|---|---|---|---|
| **Uniprot Accession Number** | **Protein names** | **Uniprot Accession Number** | **Protein name** |
| A0A210PS96 | ADP/ATP translocase | A0A8S3TBB4 | ADP/ATP translocase |
| A0A8W8N948 | Elongation factor 1-alpha | D6BJL2 | Elongation factor 1-alpha |
| A0A6J8A1B7 | EEF1A | A0A6J8A1B7 | EEF1A |
| A0AAE0SNB8 | 40S ribosomal | A0A077H0M6 | Small ribosomal subunit protein eS6 |
| A0A5J6DQS7 | Heat-shock protein 70 | A0A8B6E734 | Heat shock protein 70 kDa protein 1/8 |

Abbreviations: PD: pull-down.
